# The RelA hydrolase domain: a molecular switch for (p)ppGpp synthesis

**DOI:** 10.1101/2020.12.14.421628

**Authors:** Anurag Kumar Sinha, Kristoffer Skovbo Winther

## Abstract

Bacteria synthesize guanosine tetra- and penta phosphate (commonly referred to as (p)ppGpp) in response to environmental stresses. (p)ppGpp reprograms cell physiology and is essential for stress survival, virulence and antibiotic tolerance. Proteins of the RSH superfamily (RelA/SpoT Homologues) are ubiquitously distributed and hydrolyze or synthesize (p)ppGpp. Structural studies have suggested that the shift between hydrolysis and synthesis is governed by conformational antagonism between the two active sites in RSHs. RelA proteins of γ-proteobacteria exclusively synthesize (p)ppGpp and encode an inactive pseudo-hydrolase domain. *Escherichia coli* RelA synthesizes (p)ppGpp in response to amino acid starvation with cognate uncharged tRNA at the ribosomal A-site, however, mechanistic details to the regulation of the enzymatic activity remain elusive. Here, we show a role of the enzymatically inactive hydrolase domain in modulating the activity of the synthetase domain of RelA. Using mutagenesis screening and functional studies, we identify a loop region (residues 114-130) in the hydrolase domain, which controls the synthetase activity. We show that a synthetase-inactive loop mutant of RelA is not affected for tRNA binding, but binds the ribosome less efficiently than wildtype RelA. Our data support the model that the hydrolase domain acts as a molecular switch to regulate the synthetase activity.

## Introduction

Bacteria have evolved intricate mechanisms and responses to adapt quickly to changing and stressful environments. One of such bacterial responses is the universal stringent response. The stringent response is induced in response to amino acid starvation^1,2^, fatty acid limitation^3,4^, iron limitation^5^, heat shock^6^, glucose starvation^7^, nitrogen starvation^8^, phosphate starvation^9^ and other stress conditions^10^. The stringent response reprograms cell physiology, which facilitates stress adaptation and survival under harsh environmental conditions^11^. Furthermore, the stringent response is essential for virulence and has been shown to mediate antibiotic tolerance^12,13^. Derivatives of GDP/GTP, guanosine tetra- and pentaphosphate (collectively referred to as (p)ppGpp or alarmones), are the effector molecules of the stringent response and are synthesized/hydrolyzed by the RSH superfamily (RelA/SpoT homologues) proteins. The most commonly distributed protein of this family is the bifunctional Rel protein, which has both (p)ppGpp synthetase and hydrolase activities.

In γ-proteobacteria such as in *Escherichia coli*, the *rel* gene has been duplicated to form *relA* and *spoT*^14^.The RelA protein has only (p)ppGpp synthetase activity but carries an inactive pseudo-hydrolase domain, whereas, SpoT is a weak (p)ppGpp synthetase and exhibits strong hydrolase activity^11^. Hence SpoT is essential for cell growth unless the RelA synthetase function is compromised, as SpoT is necessary for (p)ppGpp hydrolysis^15^. Weak SpoT-dependent (p)ppGpp synthesis has been reported under multiple starvation conditions, however, RelA exclusively synthesizes (p)ppGpp in response to amino acid starvation^10,11^. These metabolic cues are not mutually exclusive and accumulating evidence suggest that diverse starvation signals including glucose- and fatty acid starvation can indirectly lead to conditions that trigger the RelA-dependent stringent response^4,7,16,17^.

RelA activation occurs when RelA binds with an uncharged tRNA at an empty A-site of a stalled ribosome, which leads to induction of (p)ppGpp synthesis^18^. Cryo-EM structures of RelA in complex with uncharged tRNA and the ribosome have revealed that the C-terminal Zinc-finger domain (ZFD) and RNA recognition motif (RRM) of RelA are responsible for ribosome binding at helix 38, the A-site finger of 23S ribosomal RNA in the 50S ribosomal subunit. The C-terminal TGS domain (ThrRS, GTPase, SpoT/RelA) is primarily involved in the recognition and binding to uncharged tRNA^19-21^ (Fig. 1A and 3A). All three domains enclose the A-site tRNA, and expose the N-terminal synthetase (SYN) and inactive hydrolase (pseudo-HD) domains on the surface of the ribosome. Recently, it was demonstrated using an *in vivo* UV crosslinking and analysis of cDNAs (CRAC) approach that RelA interacts with the ribosome as a RelA•tRNA complex^2^. RelA is thought to bind with tRNA at ribosomal A-sites during amino acid starvation, when EF-Tu•GTP•tRNA ternary complexes are scarce^18^.

**Fig. 1.**
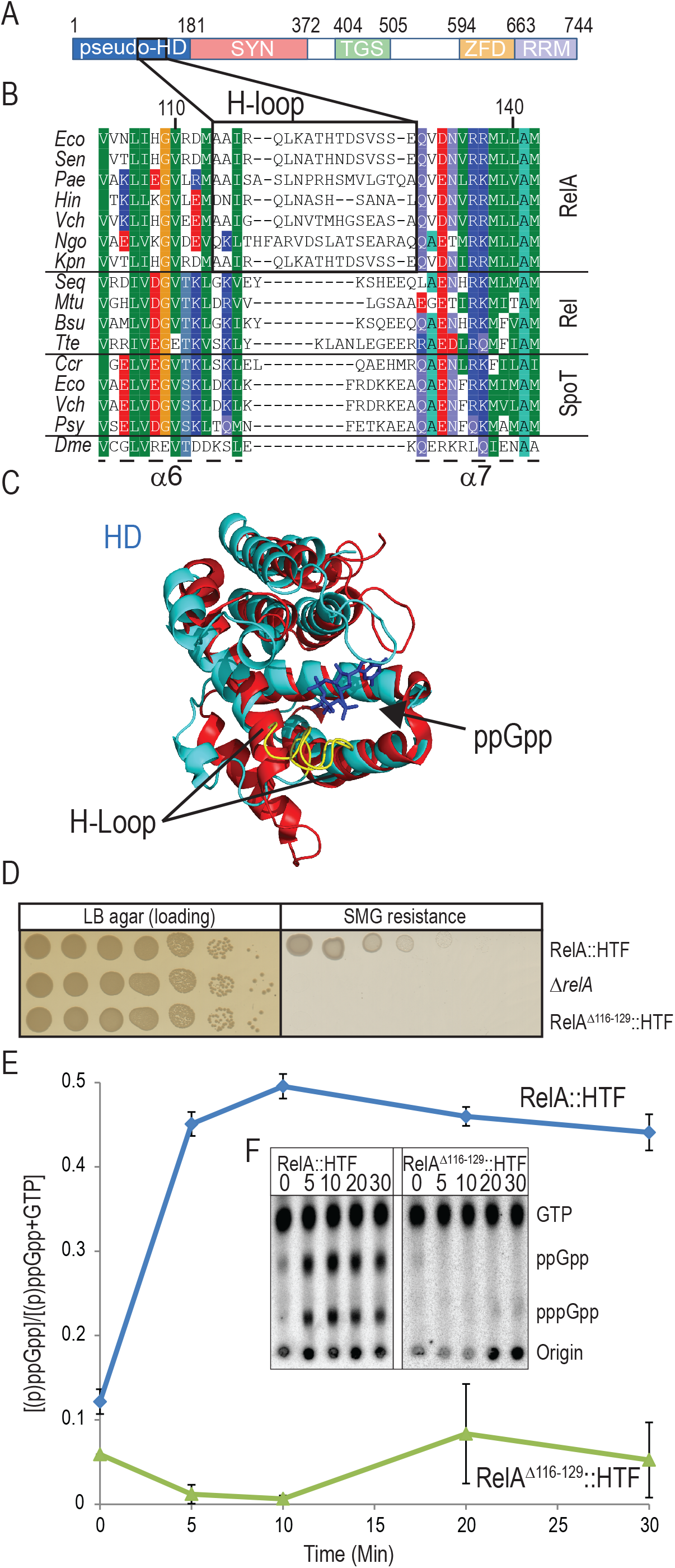
Residues of 114-130 of the RelA pseudo-hydrolase domain form a loop that controls ppGpp synthesis. **A)** Illustration of functional domains of RelA: HYD, inactive hydrolase domain (residues 1-181); SYN, synthetase domain (residues 182-372); TGS, ThrRS, GTPase and SpoT-like domain (residues 404-505); ZFD, Zinc finger domain (residues 594-663) and RRM, RNA recognition motif (residues 664-744). **B)** Multiple sequence alignment of selected RelA, Rel and SpoT sequences. *Eco, Escherichia coli, Sen, Salmonella enterica, Pae, Pseudomonas aeruginosa, Hin, Hemophilus influenza, Vch, Vibrio cholera, Ngo, Neisseria gonorrhoeae, Kpn, Klebsiella pneumoniae, Seq, Streptococcus dysgalactiae subsp. equisimilis, Mtu, Mycobacterium tuberculosis, Bsu, Bacillus subtilis,Tte, Thermus thermophilus Ccr, Caulobacter crescentus, Psy, Pseudomonas syringae, Dme, Drosophila melanogaster*. Location of helices α6, α7 and the H-loop is indicated. **C)** Structure of RelA pseudo-hydrolase domain (PDB: 5IQR, shown in cyan) superimposed onto Rel_Tte_ hydrolase domain (HD, PDB: 6S2T, shown in red). Position of ppGpp in Rel_Tte_ hydrolase active site is indicated in blue and the H-loop of RelA is shown in yellow. **D)** Functional assay of stringent response in H-loop deletion mutant RelA^Δ116-129^. *Escherichia coli* K-12 MG1655 *relA::HTF*, MG1655 *ΔrelA* and MG1655 *relA*^*Δ116-129*^::*HTF* were grown overnight in LB medium at 37°C. The cells were washed and serial diluted in phosphate buffered saline (PBS) and spotted onto LB agar (Loading) and MOPS MM SMG agar plates (SMG resistance) and plates were incubated at 30°C. **E)** and **F)** (p)ppGpp measurements in RelA H-loop mutant. Strains from D) were grown exponentially in MOPS minimal medium at 30°C containing [^32^P]-radiolabeled phosphate as described in methods. At time zero cells were starved for isoleucine by addition of 500μg/mL L-Valine (final concentration). Samples were collected at time points indicated (min), precipitated in formic acid and spotted on a TLC plate. Nucleotides were separated using 1.5M potassium phosphate pH 3.4 as solvent. E) Quantification of (p)ppGpp in four biological replicates for RelA-HTF and two for RelA^Δ116-129^::HTF with Standard deviations. F) Representative TLC from E), positions of GTP, ppGpp and pppGpp are indicated.

In bifunctional RSHs, including Rel, (p)ppGpp hydrolysis or synthesis is governed by conformational antagonism between the active sites of the hydrolase and synthase domains^22-24^. Specifically, switching between hydrolase domain ON or OFF determines if the synthetase domain will be OFF or ON. More recently, it has also been shown that the TGS domain of *Bacillus subtilis* Rel is directly involved in the repression of the synthetase domain, which keeps the enzyme in a hydrolase ON-state in the absence of deacylated tRNA and the vacant ribosomal A-site^25^.

The switch-ON signal for SpoT enzymes is still not clear, but for bifunctional Rel proteins, (p)ppGpp synthesis results from an accumulation of uncharged tRNA during amino acid starvation^26-28^. SpoT does not respond to amino acid starvation like Rel or RelA, its hydrolysis or synthesis activity is instead governed by the interaction with auxiliary protein regulators. These factors have been reported to either stimulate SpoT hydrolase or synthetase activity under diverse conditions either with high intracellular GTP levels or under fatty acid, carbon or phosphate starvation^3,29-31^. In all cases, direct interaction of regulators with specific domains of SpoT control switching of hydrolase/synthetase from ON/OFF to OFF/ON and vice versa. *E. coli* RelA contains a pseudo-HD, which is conserved but lacks the essential residues needed for (p)ppGpp hydrolysis ^14,32^ (Supplementary Fig. S1D). Previous studies suggested that an interaction between the inactive pseudo-HD of RelA and the ribosome could alter the conformation of the SYN domain to regulate RelA synthetase activity^2,20^. Interestingly, mutations in the pseudo-HD of RelA that affects (p)ppGpp synthesis have previously been isolated, which indicates that the inactive hydrolase domain has a role in the regulation of RelA synthetase activity ^33^. Moreover, degenerated inactive hydrolase domains are preserved in various RelA homologs suggesting its possible important regulatory role in RelA function^14^. However, mechanistic details of this regulation have remained elusive.

Here we identify a region in the inactive pseudo-HD (between α6 and α7), which regulates RelA synthetase activity. Interestingly, this region is extended in RelA homologs compared to Rel/SpoT and by mutagenesis of the loop we reveal several mutations that have deleterious or stimulatory effects on RelA (p)ppGpp synthetase activity. Importantly, while single-point mutant (Rel^A121E^) produced very little (p)ppGpp, ribosome interaction studies (CRAC) revealed that this mutant was still able to bind to tRNA and the ribosome, albeit less efficient to the latter. Based on these results, we propose here that the inactive hydrolase domain of RelA is important as a regulatory switch for RelA activation *in vivo*.

## Results

### The H-loop of RelA pseudo-hydrolase domain controls synthetase activity

To explore if the pseudo-HD is involved in RelA•Ribosome interaction and RelA activation, we selectively mutated residues in the hydrolase domain based on their vicinity to ribosomal RNA, when RelA is bound in the ribosomal A-site (Supplementary Fig. S1B). Alanine substitutions of these potential residues did not significantly affect RelA activity, as revealed by growth on SMG plates (Supplementary Fig. S1C). RelA is essential for growth on SMG plates, which contain high concentrations of single carbon amino acids serine, methionine and glycine and leads to isoleucine starvation^34^. We used C-terminally HTF-tagged (six histidine, TEV protease cleavage site and three FLAG epitopes) functional RelA as described previously ^2^. Mutations in the hydrolase domain have previously been reported to alter RelA (p)ppGpp synthetase activity and thereby permit deletion of the otherwise essential *spoT* locus^33^. Particularly, deletion of tryptophan 39 in α-Helix2 has previously been shown to decrease RelA activity (Supplementary Fig. S1C)^33^. This deletion is likely to result in larger structural changes in the hydrolase domain and was therefore not investigated further in our study while point mutation (W39A) does not affect the RelA activity (Supplementary Fig. S1C).

To find conserved possible regulatory regions in RelA HD domain, we aligned sequences of N-terminal domains of RelA and Rel/SpoT from different species. Interestingly, the alignment revealed a short region (Residues 114-130) in the hydrolase domain between α6 and α7 unique only to RelA homologs (Fig. 1B and Fig. S1D). This region forms an extended loop in RelA compared to Rel/SpoT and is predicted to be in the vicinity of the (p)ppGpp binding site in the hydrolase domain of Rel_Tte_ of *Thermus thermophilus*^*24*^ (Fig. 1C and Supplementary Fig. S1E). For simplicity we refer to this loop as the H-loop (Hydrolase-loop). Considering the conformational antagonism observed between the HYD and SYN domain of Rel proteins, we hypothesized that in RelA, this conserved loop could possibly regulate synthetase activity by indirectly changing the conformation of the SYN domain^22,24^.

Indeed, deletion of a part of the H-loop (Δ116-129) abolished the ability of RelA^Δ116-129^ to support growth on SMG plates (Fig. 1D and Supplementary Fig. S1F for un-tagged controls). Similar effect was also observed when assaying AT (3-amino-1,2,4-triazole) resistance which causes histidine starvation (supplementary Fig. S1G). The HTF tag allows us to compare protein expression by western blot analysis in response to amino acid starvation. The lack of growth on SMG plates cannot be explained by altered protein expression or stability, as both RelA::HTF and RelA^Δ116-129^::HTF are produced at comparable levels (Supplementary Fig. S1H, compare lane 3-4 and lane 11-12). To directly analyze synthetase activity, we measured (p)ppGpp accumulation after amino acid starvation. Consistent with the complementation results, MG1655 *relA::HTF* accumulated (p)ppGpp (∼ 5-fold) in response to isoleucine starvation whereas no significant increase in (p)ppGpp was observed in MG1655 *relA*^*Δ116-129*^::*HTF* (Fig. 1E and 1F). In conclusion, these results reveal a role of the H-loop in RelA activation.

### Single substitution mutations in the H-loop inhibit RelA synthetase activity

Deletion of a part of the H-loop (Δ116-129) can affect overall protein structure/function therefore, to investigate further the importance of the H-loop in regulating the synthetase activity, we implemented a genetic screen to isolate H-loop substitution mutants having altered ppGpp synthetase activity (Fig. 2A). Using error-prone PCR, randomized mutations were generated and inserted into the H-loop by λ-red recombination^35^. Mutants were isolated and screened on SMG plates. H-loop mutants showing growth defects were subsequently isolated and sequenced (Fig. 2B). Interestingly, a majority of substitutions (21 different, independently isolated mutants) were present in the start of the loop between residues 110-123 and had either one or two amino acid changes. Remarkably, substitutions primarily generated proline or charged amino acids such as glutamate or lysine. Four different mutants were selected from the primary screen (L119M, A121E, M113K and Q118L) and re-introduced at the *E. coli* chromosomal *relA* locus (Fig. 2C). As a control I116L was used, which has previously been isolated in the study by Montero et al. ^33^. We observed that all the tested mutations affected RelA complementation on SMG plates, however, A121E drastically hampered RelA activity (Fig. 2C). Similar results were obtained when these mutations were introduced into untagged RelA (Supplementary Fig. S2A) or when assaying AT resistance (Supplementary Fig S1G). Furthermore, comparable protein levels of RelA::HTF, RelA^A121E^::HTF and RelA^I116L^::HTF in response to amino acid starvation were confirmed by western analysis (Supplementary Fig. S1H, compare lanes 3-4 with 5-6 and 9-10). Consistent to their growth phenotype on SMG plates, MG1655 *relA*^*I116L*^::*HTF* and *relA*^*A121E*^::*HTF* showed reduced (p)ppGpp synthesis in response to isoleucine starvation as measured by thin layer chromatography (Fig. 2D, for TLCs see Supplementary Fig. S2B). Particularly, MG1655 *relA*^*A121E*^::*HTF* showed a little (p)ppGpp accumulation after 30 min (30% of wild-type after 30 min). In conclusion, we have isolated a hydrolase domain H-loop substitution mutant, RelA^A121E^ which significantly affects (p)ppGpp synthesis.

**Fig. 2.**
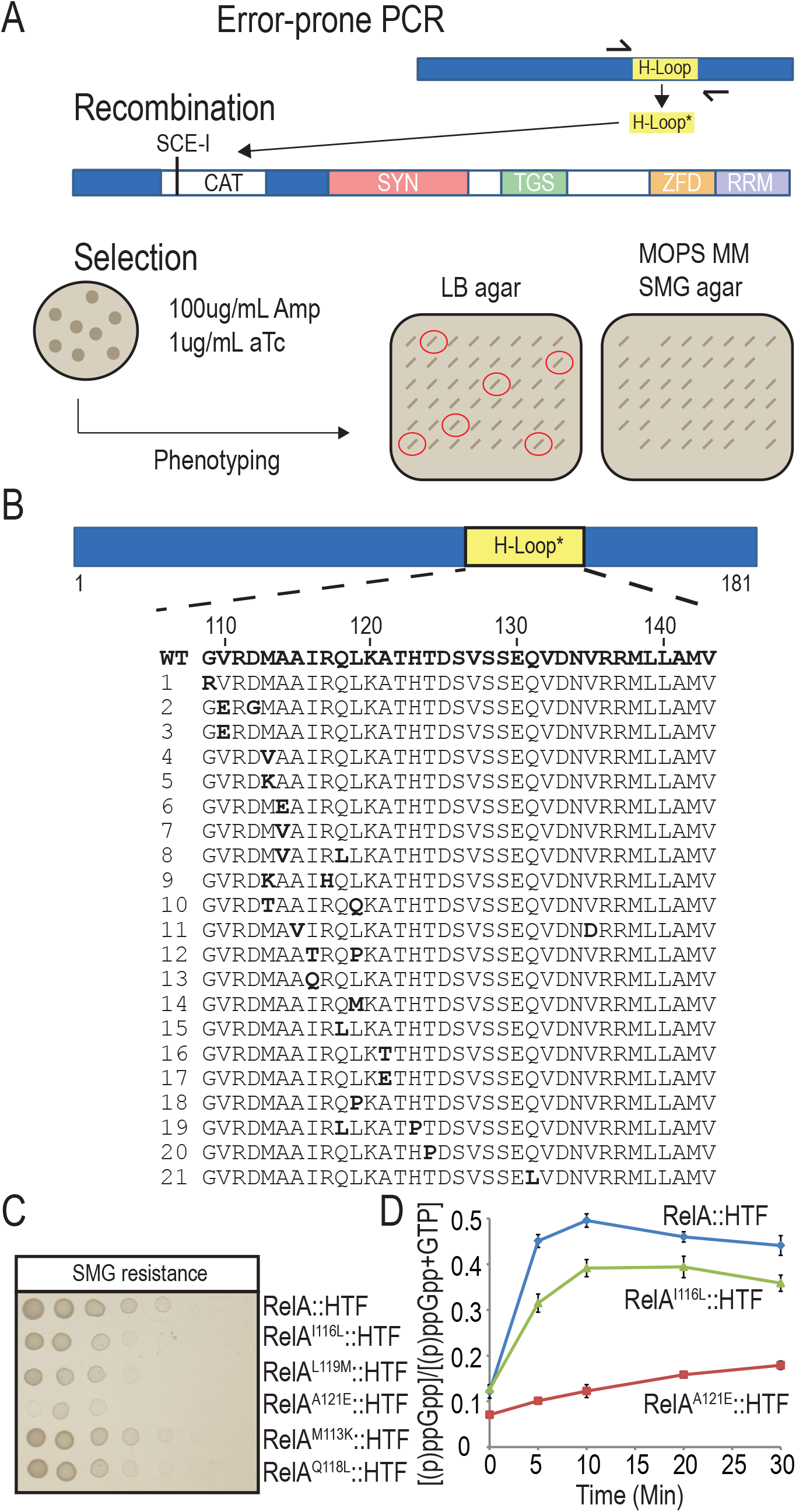
Isolation of H-loop mutants with altered (p)ppGpp synthesis. **A**) Outline of the H-loop random mutagenesis screen. The H-loop was amplified using error-prone PCR as described in methods. The PCR product was then electroporated into MG1655 *relA*^*I116::cm*^::*HTF* containing plasmid pWRG99 which have previously been induced with 0.2% arabinose to express lambda recombinase^35^. After phenotypic expression at 37°C, cells were plated on LB agar containing 100μg/mL ampicillin (Amp) and 1μg/mL anhydrotetracycline (aTc). Induction of the *Sce-I* restriction enzyme by aTc addition facilitated the site-directed replacement of the chloramphenicol cassette (CAT) with the PCR product. Colonies were selected and re-streaked onto LB agar (loading control) and MOPS MMl SMG plates to assay RelA functionality at 30°C. Colonies that showed decreased growth on functional plates were sequenced (indicated with red circles). **B**) Overview of the H-loop and substitution mutants isolated in A). Repeated mutations, nonsense and frame-shift mutations were excluded in this study. **C**) Assaying the stringent response in selected substitution mutants. Substitution mutations I116L, L119M, A121E, M113K and Q118L were introduced by site-directed recombination in MG1655 *relA::HTF* as described in methods. The cells were grown in LB 37°C, washed in PBS and spotted onto MOPS MM SMG plates (SMG resistance) followed by incubation at 30°C. **D**) (p)ppGpp measurements in selected mutants in response to isoleucine starvation. MG1655 *relA::HTF, relA*^*I116L*^::*HTF* and relA^A121E^::HTF were grown exponentially in MOPS minimal medium containing ^32^P-labeled phosphate. To induce isoleucine starvation, L-Valine was added, to a final concentration of 500μg/mL. Samples were collected before (time zero) and after starvation, followed by precipitation and separation by thin layer chromatography. The curves from mutants are based on two biological replicates with standard deviations.

### RelA^A121E^ substitution mutant interacts with uncharged tRNA and the ribosome

We have implemented a crosslinking methodology, crosslinking and analysis of cDNA (CRAC) (Supplementary Fig. S3A), which allows mapping of protein-RNA interactions with single nucleotide resolution, in living cells ^2,36^. We applied this approach to investigate the interaction of the H-loop RelA mutant with the tRNA and the ribosome. In response to isoleucine starvation, wild-type RelA crosslinking increases with uncharged isoleucine tRNA, tRNA^ileTUV^, and the ribosome ^2^. Using false discovery rate analysis (FDR) we previously identified three sites on the ribosome, which showed statistically increased crosslinking with RelA after isoleucine starvation. These sites are: the A-site finger (ASF) of 23S rRNA, Helix 15 (H15) of 16S rRNA and the Sarcin-Ricin Loop (SRL) of 23S rRNA (Fig. 3A). Consistent with the cryo-EM structure of RelA bound to the ribosome along with the uncharged tRNA^19-21^, we observed crosslinking between the ZFD and RRM domains of RelA with the ASF of 23S rRNA, and the TGS domain of RelA with H15 of 16S rRNA ^2^. These interactions are consistent with the RelA accommodation in the A-site during isoleucine starvation. Here we performed CRAC to compare interactions of the RelA::HTF and RelA^A121E^::HTF with the tRNA and the ribosome.

**Fig. 3.**
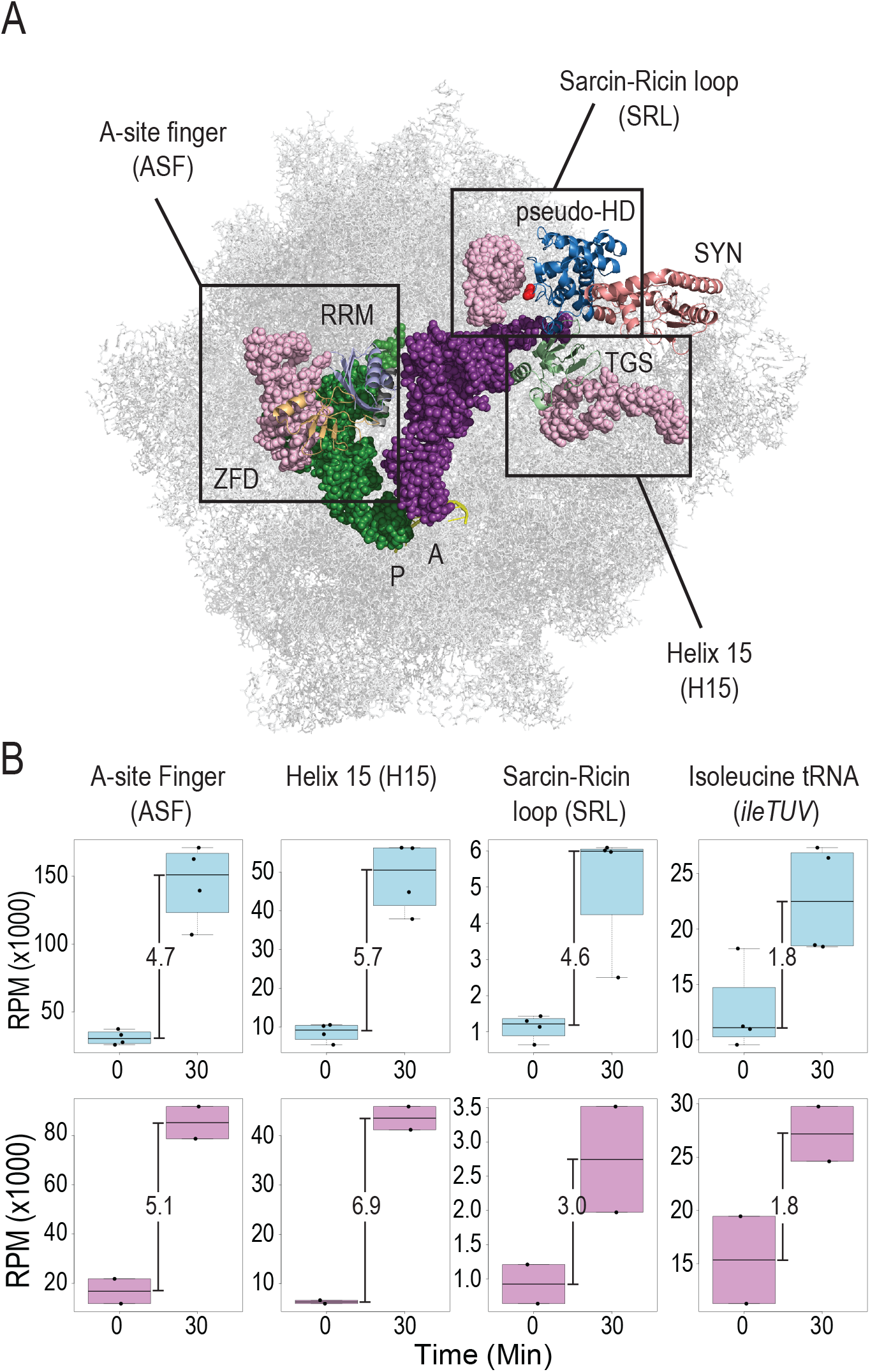
RelA^A121E^ displays decreased interaction with the ribosome in response to isoleucine starvation. **A**) Structure of RelA bound to tRNA and the ribosome (PDB: 5IQR). Functional domains have been colored as in Fig. 1A) and A- and P-site tRNAs are indicated in magenta and green respectively. Previously identified RelA ribosome interaction sites using false discovery rate (FDR) analysis ^2^: A-site Finger (ASF, nucleotide 834-927 in 23S rRNA), Sarcin-Ricin Loop (SRL, nucleotide 2652-2673 in 23S rRNA) of 50S ribosomal subunit and Helix 15 (H15, nucleotide 328-407 in 16S rRNA) of 30S ribosomal subunit are indicated with boxes in light pink. Location of A121 in the hydrolase domain is indicated in red **B**) Boxplots of Normalized cDNA coverage with ribosome and tRNA obtained from CRAC analysis of MG1655 *relA::HTF* (cyan) and MG1655 relA^A121E^::HTF (pink) before (0) and 30 minutes after isoleucine starvation (30). Reads from biological replicates are shown as dots are derived from overlaps with interaction sites indicated in A) and normalized to reads per million (RPM). Fold-change between unstarved (0) and starved (30) cells are indicated.

The main interaction site for RelA C-terminal ZFD and RRM domains is helix 38 of 23S rRNA A-site finger (ASF), which bridges the A-site between the ribosomal subunits ^2,19,20,37^ (Fig. 3A). RelA and RelA^A121E^ showed similar fold increase in interaction with ASF after isoleucine starvation, 4.7- and 5.1-fold respectively (Fig. 3B and Supplementary Fig. S3B and S3E).

However, the number of normalized cDNA reads in RelA^A121E^ is only 59% of the wild type, which indicates weaker binding to the ribosome. RelA^A121E^ also showed weaker interaction with the Sarcin-Ricin Loop (SRL) of 23S rRNA i.e. 53% of the wild type and lower fold increase in response to starvation (3-fold compared to 4.6-fold) indicating decreased interaction at this site by the mutant RelA protein (Fig. 3B and Supplementary Fig. S3C). Interestingly, the interaction of RelA^A121E^ TGS domain with Helix 15 (H15) of 16S rRNA was less affected and only showed 10% lower number normalized cDNA reads after 30 minutes of amino acid starvation (Fig. 3B and Supplementary Fig. S3C and S3F). The TGS interaction occurs with the ribosome when RelA binds with the uncharged tRNA in the ribosomal A-site^21^. We did not observe any significant difference in the binding with uncharged isoleucine tRNA (tRNA^IleTUV^) between RelA::HTF and RelA^A121E^::HTF in response to isoleucine starvation (Fig. 3B and Supplementary Fig. S3D and S3G). This suggests that the mutant protein binds efficiently to tRNA, similar to wild-type protein. In RelA::HTF, (p)ppGpp accumulated within 5 minutes of isoleucine starvation and we therefore performed CRAC also at this time-point (Supplementary Fig. S3E-G). However, we did not observe any significant difference in the enrichment patterns of the ribosomal RNAs (23S and 16S) and isoleucine tRNA between 5 min and 30 min of starvation in both the strains. Based on these results, we conclude that RelA^A121E^::HTF is still able to bind to uncharged tRNA and the ribosome in response to isoleucine starvation. However, we did observe a lower number of cDNA reads from the ribosomal RNA suggesting a weaker binding to the ribosome and in particular a weaker interaction with the ASF and SRL in the mutant. The results indicate that the intact pseudo-HD domain of RelA is necessary for efficient ribosome binding and is important for (p)ppGpp synthesis.

### RelA^A121E^ crosslinks to the same RNA sites as RelA

After confirmation that both RelA::HTF and RelA^A121E^::HTF bind to similar sites on the ribosome, we analyzed the crosslinking pattern at the specific residue level. In CRAC method, crosslinking pattern can easily be scored as reverse transcription mutations (RT-mutations) that occur at high frequency at the crosslinking sites. The most common mutations observed are deletions or substitutions in the cDNA at the crosslinking sites^2,38^. Crosslink mediated increase in both deletions and substitutions were clearly observed in the ASF region of 23S rRNA in response to isoleucine starvation (Supplementary Fig. S4A-D for all RT-mutation heat maps). We primarily observed substitutions at A887 and deletions at U884-C888 showing crosslinking pattern of the ZFD domain (Fig. 4A and 4B). These mutations were similar for both RelA and RelA^A121E^, suggesting similar binding of the ZFD domain to the ASF. The signature of RRM domain interactions with ASF is deletions at position U894-A896 (Fig. 4A and 4B)^2^. Though the number of cDNA reads from RelA^A121E^ aligning to ASF crosslinking is lower than the RelA, the crosslinking patterns were the same. Furthermore, crosslinking of the TGS domain to Helix 15 of 16S rRNA resulted in significant substitutions at U368 and deletions at position G359, A364, U365 and A367 (Fig. 4C and 4D and Supplementary Fig. S4E-H). At Helix 15, we observed slightly lower crosslinking with RelA^A121E^ as compared to RelA, but with similar crosslinking pattern. Another crosslinking site in the ribosome is the SRL of 23S rRNA, which showed increased amounts of substitutions at A2660 and deletions at U2653-G2655, G2659 and A2660 (Fig. 4E and 4F and Supplementary Fig. S4I-L). Again, the crosslinking pattern is the same between the two strains, but the number of reads was lower in RelA^A121E^. Additionally, as expected from the enrichment data, crosslinking to tRNA^ileTUV^ was similar for both RelA and RelA^A121E^ (Supplementary Fig. S4N-M). Taken together, the crosslinking data indicate that RelA and RelA^A121E^ bind to uncharged tRNA and to the ribosome in a similar manner but mutant protein interacts less efficiently with the ribosome compared to wild-type protein. Importantly, the interactions with ASF and SRL are affected significantly indicating that these interactions might play an important role for SYN domain activation and (p)ppGpp synthesis.

**Fig. 4.**
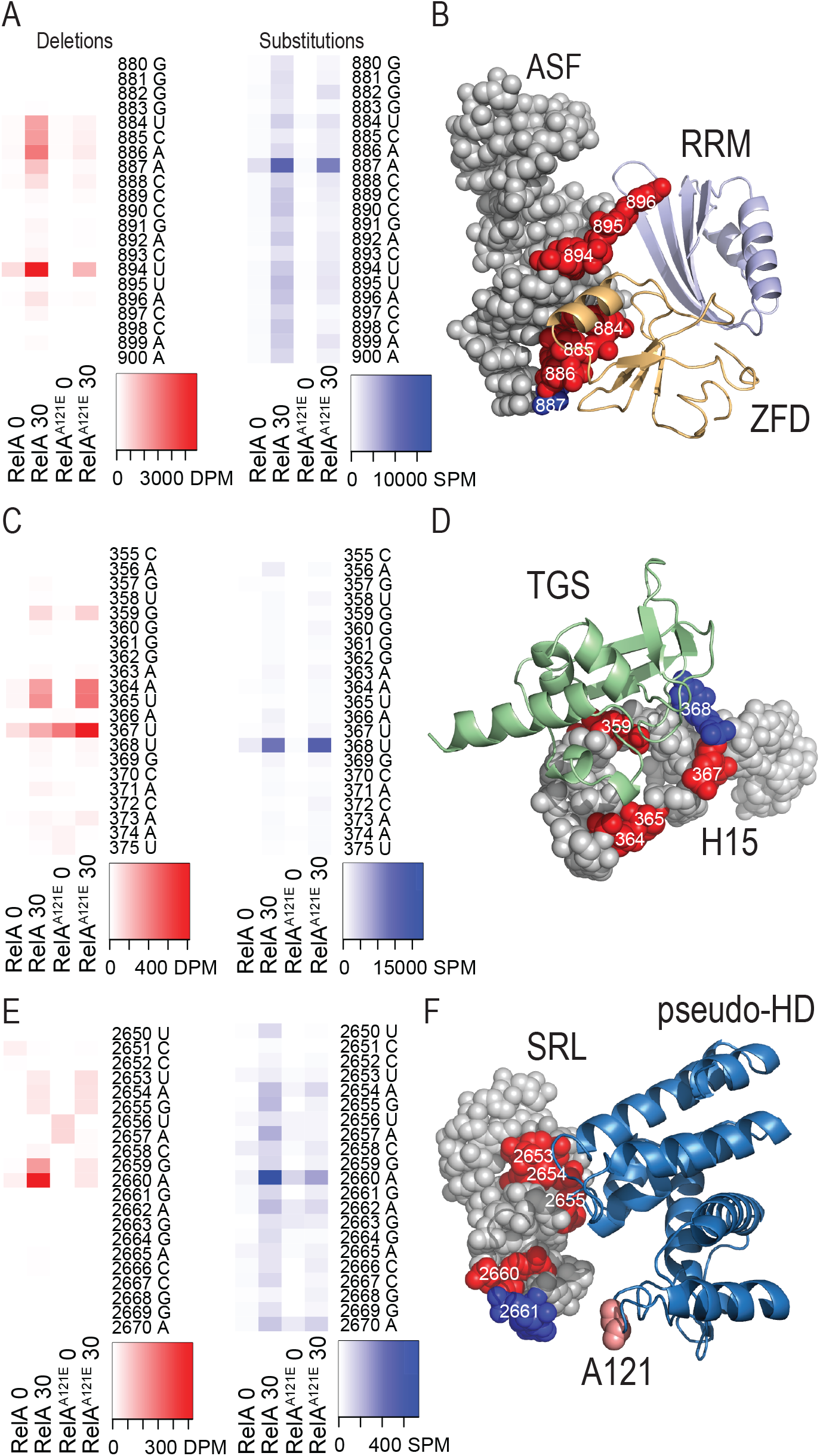
RelA^A121E^ ribosome interaction is analogous to RelA. RT-mutations (deletion and substitutions) in cDNA reads obtained with CRAC analysis of MG1655 encoding *relA::HTF* or *relA*^*A121E*^::*HTF* before and 30 min after isoleucine starvation (here named 0 or 30, respectively). Heat maps show nucleobase positions with increased error-frequencies caused by RelA•RNA crosslinking. Deletions per million, DPM, or Substitutions per million, SPM, are indicated in red and blue respectively. **A**) RT-mutations in the A-site finger (ASF) of 23S rRNA (nt 880-900). **B**) Close-up on the ASF (PDB: 5IQR) with positions with significant number of RT-mutations. Deletions or substitutions are shown in red and blue, respectively. ZFD and RRM domains of RelA are colored in pale orange and pale blue. **C**) RT-mutations in helix 15 (H15) of 16S rRNA (nt 355-375). **D**) Close-up on H15 (PDB: 5IQR) with crosslinking sites. TGS domain of RelA is displayed in pale green. **E**) RT-mutations in the Sarcin-Ricin Loop (SRL) of 23S rRNA (nt 2650-2670). **F**) Close-up on SRL (PDB: 5IQR) with crosslinking sites. The pseudo-HD of RelA is shown in blue and the position of mutated alanine 121 is highlighted in salmon.

### Alanine substitutions in the H-loop stimulate (p)ppGpp synthesis

The data presented above provided evidence that the H-loop of the pseudo-HD is important for the regulation of the SYN domain and (p)ppGpp synthesis. Moreover, single or double substitutions in the H-loop decreased ribosome binding and affected for (p)ppGpp synthesis. To investigate further the role of the H-loop, we made site-directed substitutions of four residues of the H-loop into alanine, which would significantly affect the structure of the loop. The mutated residues were arginine 117, glutamine 118, lysine 120 and histidine 123 (Mutant is referred to as RelA^QUAD^::HTF,Fig. 5A).

**Fig. 5.**
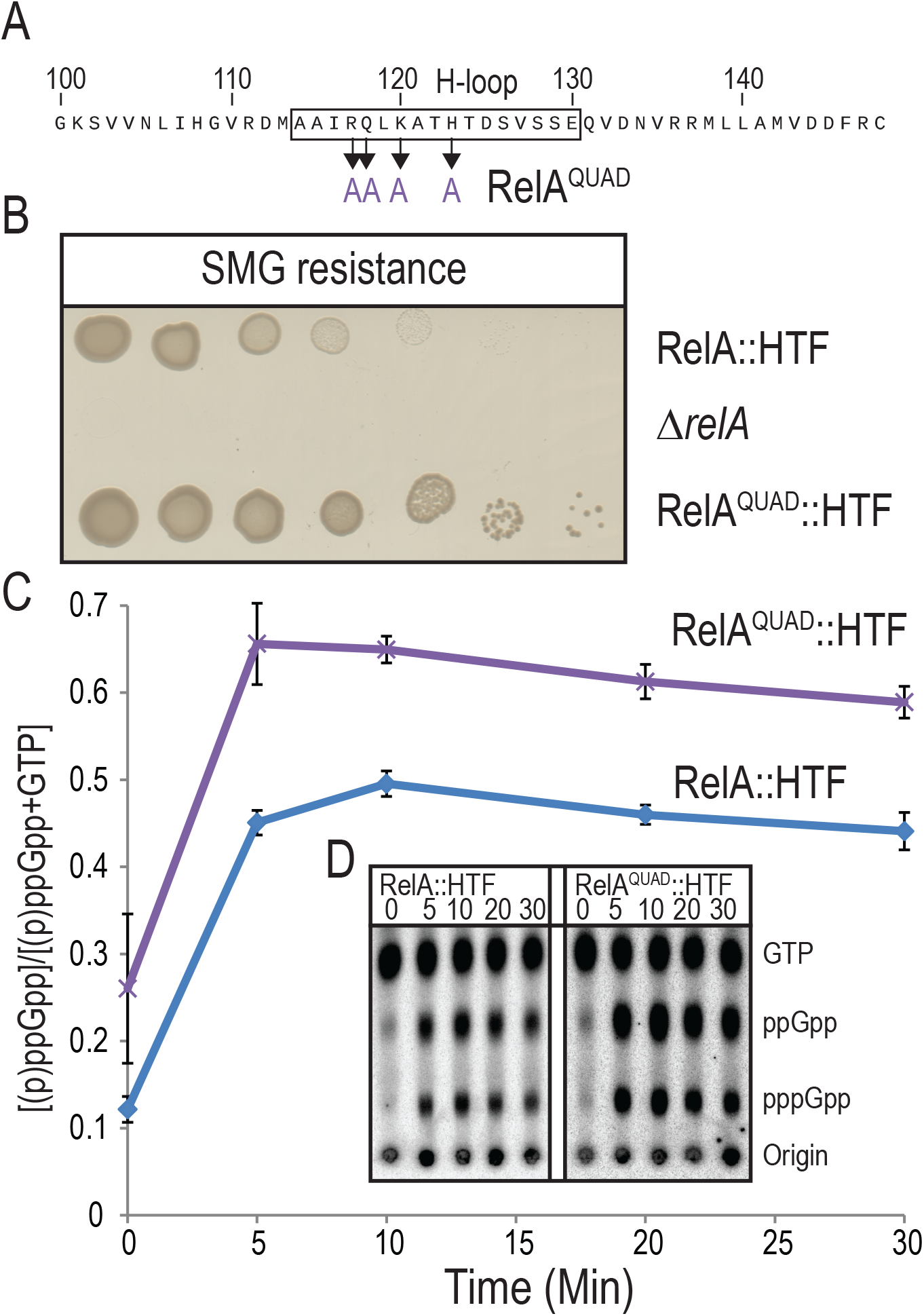
Alanine substitutions in the H-loop leads to increased (p)ppGpp synthesis. **A**) Primary sequence of the H-loop region (indicated within a box) of RelA. Arrows indicate positions which have been substituted to alanine in RelA^QUAD^ (R117A, Q118A, K120A and H123A). **B**) Functional assay of RelA^QUAD^. Cell cultures of MG1655 *relA::HTF*, MG1655 *ΔrelA* and MG1655 *relA*^*QUAD*^::*HTF*, were washed in PBS, serial diluted and spotted onto MOPS MM SMG plates (SMG resistance). The plates were incubated ON at 30°C. **C)** (p)ppGpp measurements of strains in A). Cells were grown exponentially in MOPS minimal medium containing ^32^P-labeled phosphate. Isoleucine starvation was induced by addition of L-valine, to a final concentration of 500μg/mL. Samples were collected before (time zero) and after starvation, precipitated and separated by thin layer chromatography. The RelA^QUAD^-HTF curve is based on two biological replicates with standard deviations. **D)** Representative TLC from C), positions of GTP, ppGpp and pppGpp are indicated.

Surprisingly, the mutant showed increased survival on SMG plates as compared to wild-type (Fig. 5B). The stimulatory effect of the RelA^QUAD^ mutant was independent of the HTF tag, as a similar effect was observed with the untagged protein (Supplementary Fig. S1F). A similar effect was also observed when assaying AT resistance (Supplementary Fig. S1G) In an attempt to explain this surprising effect, (p)ppGpp synthesis was measured by TLC after isoleucine starvation (Fig. 5C and 5D). Interestingly, while RelA showed an approximately 5-fold increase in (p)ppGpp synthesis, RelA^QUAD^::HTF only showed about a 3-fold increase in response to isoleucine starvation. The strain expressing RelA^QUAD^::HTF on the other hand, already had an elevated basal level of (p)ppGpp prior to starvation (about 2-fold higher than the basal level of RelA). The RelA^QUAD^::HTF protein levels were comparable to wild-type RelA before and after starvation and hence the higher basal level of (p)ppGpp is due to the higher basal synthetase activity of the mutant protein rather than the amount of protein *per se*. (compare lanes 7-8, Supplementary Fig. S1H). This suggests that the RelA^QUAD^ mutant intrinsically produces more (p)ppGpp than the wildtype RelA. This feature surprisingly did not significantly affect growth on LB plates, but supported better survival and growth on SMG and AT plates. In conclusion, four alanine substitutions in the H-loop of RelA increased basal level of (p)ppGpp synthesis, but decreased the fold induction of (p)ppGpp synthesis in response to isoleucine starvation. The increased (p)ppGpp synthesis by the RelA^QUAD^ mutant even in the absence of a starvation signal further demonstrates the importance of the H-loop in regulation of the SYN domain of RelA.

## Discussion

Here, we have established a new fundamental function of the N-terminal inactive pseudo-hydrolase domain (pseudo-HD) of RelA for the activation of the synthetase domain (SYN). By genetic, biochemical and *in vivo* crosslinking analysis, we show that the HYD domain is important for regulating RelA synthetase activity. Further, we identify a loop between helix α6 and α7 (Residues 114-130), which is divergent and longer in hydrolase inactive RelA homologues as compared to hydrolase active Rel/SpoT (Fig. 1B). Random mutagenesis in this loop led to the identification of residues, which when mutated, affects RelA synthetase activity. Introduction of multiple alanine substitutions in the loop (RelA^QUAD^) has an stimulatory effect on (p)ppGpp synthesis and increases the basal level before isoleucine starvation (Fig. 5). Surprisingly, a single point mutant RelA^A121E^ in the pseudo-HD severely affects SYN domain function and (p)ppGpp synthesis (Fig. 2).

Recently, it has been shown in the Rel_Tte_ of *Thermus thermophiles*, that the loop between α6 and α7 changes conformation and moves away from hydrolase active site to make hydrolase active site free for ppGpp hydrolysis and goes towards it to block the hydrolase active site^24^. This conformational change is mediated by binding of ppGpp to the hydrolase active site of the protein. Additionally, binding of ppGpp to the hydrolase domain directly precludes the binding of GDP/ATP in the synthetase domain to prevent ppGpp hydrolysis and synthesis happening simultaneously^24^. Similar observations had also been reported for Rel_seq_ of *Streptococcus dysgalactiae*, here the binding of pGpp in the hydrolase domain does not prevent binding of GTP or GDP in the synthetase domain, but prevents binding of ATP and (p)ppGpp synthesis^22,23^. Recently it was shown that the TGS domain of Rel_Bsu_ is involved in the regulation of the synthetase domain^25^. Upon binding of Rel_Bsu_ to the ribosome along with uncharged tRNA, the TGS domain moves away from the synthetase domain with structural changes in the N-terminal domains including α7 of the hydrolase domain. Moreover, substitutions in α6 and α7 (R125E/M127) were observed to decrease hydrolase activity and increase synthetase activity^25^.Thus, in all cases the hydrolase domain seems to control (p)ppGpp synthesis.

In *E. coli* RelA, the hydrolase domain is inactive and crucial residues for hydrolytic function including the conserved HDxxED motif is absent (Supplementary Fig. S1D). However, our results show that the hydrolase domain still plays a similar role of regulating SYN domain function and that might be the reason for it to be preserved in the RelA even with the extended H-loop between helix α6 and α7.

The single point mutant RelA^A121E^ identified in our study is severely affected for (p)ppGpp synthesis (Fig. 2). This mutant was still able to bind to the ribosome and uncharged tRNA in response to amino acid starvation, which is consistent with the fact that the substitution is present in the pseudo-HD, distant from ribosome and tRNA binding domains (Fig. 3). The similar binding pattern with the ribosome and tRNA of this mutant confirms that the mutant protein is structurally similar to wild-type RelA protein and this single point mutation has not affected protein structure drastically. Interestingly, while crosslinking to Helix 15 of 16S rRNA is similar between RelA and RelA^A121E^, crosslinking to the ASF and the SRL is slightly affected (59% and 53%, respectively, Fig. 3). Thus far, we know that the ZFD and RRM domains are responsible for the binding to ASF however; the mechanistic details of RelA crosslinking to SRL is still not clear^2,19,20,37^. Due to the location of the SRL in the vicinity of the pseudo-HD and the dynamic nature of the N-terminus, we earlier suggested that the pseudo-HD interacts with SRL and could promote RelA activation^2^. Though crosslinking with the SRL was found to be lower in the RelA^A121E^, the crosslinking pattern is similar to wild-type suggesting that the H-loop is perhaps not responsible for this interaction (Fig. 4). Previously, it has been shown that cleavage between G2661 and A2662 of SRL by α-sarcin toxin decreases GTPase activity of EF-G when bound to the ribosome, but does not affect RelA (p)ppGpp synthesis *in vitro* and argued that SRL is not needed for RelA acitvity^37^. In addition interaction with the SRL was also not observed by Rel_Bsu_ from *Bacillus subtilis*^*25*^.

Nevertheless, it has previously been shown that ribosomal protein L11 and the SRL contacts with the RelA bound tRNA at the elbow and acceptor stem to stabilize the distorted configuration (Supplementary Fig. S1A)^20^. Therefore, the RelA-SRL interaction might be dynamic and occur during binding of RelA to the ribosome to help RelA to be stabilized on the ribosome.

We argue that the most probable explanation is that the RelA^A121E^ mutant is locked in a conformation on the ribosome that does not allow the SYN domain to have sufficient flexibility to reach a fully active configuration. It should be noted that RelA^A121E^ is not a null mutant and it can still synthesize ppGpp but with reduced efficiency (Fig. 2D). Another possibility is that the (p)ppGpp synthesis could stabilize the binding of RelA to the A-site and promote increased synthesis by a mechanism similar to positive allosteric feed-back^37,39,40^. In either case, RelA needs the fully functional H-loop to bind efficiently to the ribosome and to switch ON (p)ppGpp synthetase activity of the SYN domain.

In conclusion, we present robust evidence demonstrating that the inactive HYD domain of RelA plays a novel regulatory role in controlling (p)ppGpp synthetase activity of the SYN domain. Our data thus unravel a distinct layer of RelA synthetase activity regulation. We believe that the ON/OFF switch earlier proposed for Rel proteins might be a well-conserved regulatory mechanism for RSH proteins, which has evolved differently for monofunctional and bifunctional proteins.

## Materials and Methods

### Strains and plasmids

Strains constructed and plasmids used in this study are described in supplementary information. Strains and oligonucleotides are listed in Supplementary table S1.

### Media and growth conditions

*Escherichia coli* K-12 strains were routinely grown in liquid LB complex medium or on solid LB agar medium at 30 or 37°C. For amino acid starvation experiments the bacterial cells were grown in MOPS (morpholinepropanesulfonic acid) minimal medium at 30°C or 37°C supplemented with 0.2% glucose, with all nucleobases (10 µg ml^-1^ of each) and 1.32mM K_2_HPO_4_ ^41^. In liquid medium, isoleucine starvation was induced by addition of L-Valine to a final concentration of 500μg/mL^42^. For functional studies on solid medium, isoleucine starvation was induced by the addition of single carbon amino acids: Serine, Methionine and Glycine (SMG), to a final concentration of 100μg/mL on MOPS minimal medium (MM) agar plates^34^. In addition, RelA functionality was also assayed by 3-amino-1,2,4-triazole (AT) resistance^43^. AT resistance was assayed on M9 minimal medium agar containing 0.2% glucose, all amino acids except histidine, 1mM adenine, 1mM thiamine with or without 15mM 3-amino-1,2,4-triazole. To select for resistance cassette on chromosome and plasmid, liquid or solid media were supplemented with 10/25μg/mL chloramphenicol or 100μg/mL ampicillin. When stated, 0.2% arabinose or 1μg/mL anhydrotetracycline was added to induce transcription from arabinose or tetracycline inducible promoters.

### (p)ppGpp measurements

(p)ppGpp measurements during isoleucine starvation was performed as described previously^41^. Overnight cultures of relevant strains were diluted 100-fold in 5 ml of MOPS minimal medium supplemented with 0.2% glucose and all nucleobases (10 µg ml^-1^ of each), and incubated at 30°C with shaking. At OD_600_ ∼0.5, cells were diluted 10-fold to an OD_600_ of ∼0.05 and were left to grow with shaking at 30°C with H_3_^32^PO_4_ (100 µCi/ml). After ∼2 generations (OD_600_ of ∼0.2), amino acid starvation was induced by the addition of valine (500 μg/ml). Fifty-microliter samples were withdrawn before and 5, 10, 20 and 30 min after addition of valine. The reactions were stopped by the addition of 10 μl of ice-cold 2 M formic acid and centrifuged at maximum speed for 1hr at 4°C. 10 μl of each reaction mixture was loaded on polyethyleneimine (PEI) cellulose thin-layer chromatography (TLC) plates (purchased from GE Healthcare) and separated by chromatography in 1.5M potassium phosphate at pH 3.4. The TLC plates were revealed by phosphorimaging (GE Healthcare) and analysed using ImageQuant software (GE Healthcare). The increase in the level of (p)ppGpp was quantified as the fraction of (p)ppGpp of (p)ppGpp+GTP.

### Random mutagenesis screening using error-prone PCR

To screen for RelA hydrolase mutants with altered synthetase activity the loop region was amplified from *relA* using oligos loop-mut-f and loop-mut-rv using the DreamTaq polymerase (Thermo) According to ^44^. 100 μl PCR was prepared containing 10 U DreamTaq polymerase, 10 μl 10X Dream tag buffer, 200 μM of each dNTP, 0.3 μM of each primer, colony DNA as template and 2-4 μl mutagenesis buffer (4 mM dTTP, 4 mM dCTP, 2.5 mM MnCl_2_, 27.5 mM MgCl_2_). The PCR product was purified and electroporated into recombination competent MG1655 *relA*^*I116::cm*^::*HTF* containing plasmid pWRG99. After 1hr of phenotypic expression cells were serially diluted and plated on LB-plates containing 100μg/mL ampicillin and 1μg/mL anhydrotetracycline. Positive Sce-I resistant clones were re-streaked on LB plates and MOPS MM SMG plates at 30°C.

### RelA-RNA interactions by UV crosslinking and analysis of cDNAs

Crosslinking and analysis of cDNAs (CRAC) was performed essentially as previously described in ^2^ (Fig. S3A shows an overview). MG1655 *relA::HTF* and MG1655 *relA*^*A121E*^::*HTF* were grown overnight (ON) MOPS minimal medium supplemented with 0.2% glucose and all nucleobases (10 µg ml^-1^ of each) at 30°C. The ON cultures where then diluted to OD_600_ = 0.005 into two flasks containing 2L MOPS minimal medium and incubated with shaking at 30°C. At OD_600_ = 0.2 one culture was UV crosslinked in a W5 crosslinking unit (Van Remmen UV techniek) by irradiation with 1800 mJ of UV-C for 100 seconds. The other culture was starved for Isoleucine by addition of 500μg/mL L-Valine for 5 or 30 min before exposure to UV. After UV irradiation the cultures were separated into 1L aliquots, harvested and the pellet washed in ice-cold 1XPBS (Phosphate Buffered Saline, Oxoid) before rapid freezing in liquid nitrogen. The pellets were stored at −80°C before proceeding with purification. The HTF-tagged proteins were then purified, linkers ligated to crosslinked RNA, cDNA synthesized and DNA libraries generated as described in^2^. The pellets were dissolved 1mL Lysis buffer (50mM Tris-HCl pH7.8, 150mM NaCl, 0.1% NP-40, 5mM β-Mercaptoethanol and Complete protease inhibitor) and lysed by vortexing 5×1 min with 3mL 0.5mmm Zirconia beads (Thistle Scientific). Lysates were cleared by centrifugation and incubated with 200μL anti-FLAG M2 affinity gel (Sigma-Aldrich) for 2 hrs at 4°C. The resin was washed twice with Wash buffer (50mM Tris-HCl pH7.8, 0.1% NP-40, 5mM β-Mercaptoethanol and 1M (high salt) or 150mM NaCl (low salt), respectively. The resin was resuspend in 600μL low salt Wash buffer and RelA was cleaved from the resin by treatment with 5μL HaloTEV protease (promega) for 2hrs at 18°C. Crosslinked RNA in the cleaved sample (500μL) was trimmed using 1μL (0.7U) RNaseIT (Agilent Technologies) for 5min at 37°C and stopped by addition of guanidine-HCl to a final concentration of 6M. The trimmed sample was subsequently bound to 100μL Ni-NTA superflow agarose (QIAGEN) over night at 4°C in Denaturing buffer (50mM Tris-HCl pH7.8, 300mM NaCl, 6M guanidine-HCl, 0.1% NP-40, 5mM β-Mercaptoethanol) with 10mM Imidazole. The resin was then washed twice in Denaturing buffer and three times in Reaction buffer (50mM Tris-HCl pH7.8, 10mM MgCl_2_ and 5mM β-Mercaptoethanol supplemented with 0.5% NP-40. First the crosslinked RNA was dephosphorylated using 0.1U/μL FastAP (Thermofischer) in Reaction buffer containing 1U/μL RNAsin (Promega) for 45min at 37°C and stopped by washing once with Denaturing buffer and three times with Reaction buffer supplemented with 0.5% NP-40. 1mM of 3’-end mirCat-33 linker (see Table S1) was ligated to the RNA using T4 RNA Ligase I (New England Biolabs) in Reaction buffer containing 1U/μL RNAsin for 6hrs at 25°C. The reaction was stopped by washing once with Denaturing buffer and three times with Reaction buffer supplemented with 0.5% NP-40. The RNA was then 5’-end phosphorylated using T4 polynucleotide kinase (Thermofischer) and 0.5μCi/μL [γP^32^]-ATP for 40min in Reaction buffer at 37°C. ATP was added to a final concentration of 1.25mM and incubation was continued for 20 min. The reaction was stopped by washing once in Denaturing buffer and three times with Reaction buffer supplemented with 0.5% NP-40. Barcoded 5’-linker (1.25mM of L5Aa, L5Ab, L5Ad, L5Bb, L5Bc or L5Bd in Table S1) was ligated to the sample RNA using T4 RNA ligase I in Reaction buffer containing 1U/μL RNAsin over night at 16°C. The resin washed three times in Wash buffer (50mM Tris-HCl pH7.8, 50mM NaCl, 0.1% NP-40 and 5mM β-Mercaptoethanol) and the RelA-RNA complex eluted twice with 200μL Elution buffer (50mM Tris-HCl pH7.8, 50mM NaCl, 150mM Imidazole, 0.1% NP-40 and 5mM β-Mercaptoethanol). The eluate was then precipitated using Trichloroacetic acid and the precipitate dissolved in 1xLDS loading buffer (Life technologies) before separation on 4-12% NuPAGE gradient gel (Life technologies) in 1xMOPS running buffer (Life technologies). RelA-RNA complexes were transferred to a Hybond C+ extra membrane (Amersham) and extracted by incubation with 100mg Proteinase K (Thermofischer) in 400 μL Wash buffer containing 1% SDS and 5mM EDTA for 2hrs at 55°C. The RNA was isolated by phenol:chloroform:isoamylalcohol and chloroform extraction followed by ethanol precipitation at -80°C for 30min. The RNA was the converted to cDNA using Superscript III reverse transcriptase (Invitrogen) and 1mM 33-rev oligo (see Table S1) at 50°C for 1hr followed by incubation with 0.5U/μL T4 RNase H at 37°C.The libraries were generated by PCR using LA Takara taq polymerase (Clontech) and oligos P5 and PE (see Table S1) size selected on a agarose gel and extracted using the MINelute extraction kit (QIAGEN).

The DNA libraries were sequenced on the Illumina MiSeq platform (50bp single-end reads) and the sequencing output analysed using the pyCRAC software package^45^. We have previously adapted this approach for RelA ^2^. FastQ files were demultiplexed using pyBacodeFilter.py and the reads 3’-end trimmed using the cutadapt tool. The reads were then collapsed based on the read sequence and the random triplet sequence in the 5’-end linker using pyFastDuplicateRemover.py. The cDNA reads were aligned to reference genome (MG1655 *E. coli* K-12 NC_000913.3) using Bowtie 2. To eliminate the possibility of alignment to identical or highly similar rRNA and tRNA we masked these in the reference genome (*alaX, alaU, alaV, argY argZ, argQ, asnU, asnV, asnW, aspU, aspV, glnW, gltU, gltV, gltW, glyW, glyX, glyY, ileU, ileV, leuQ, leuV, leuP, lysQ, lysV, lysW, lysY, lysZ, metW, metZ, metY, pheV, serX, tyrU, tyrV, valW, valU, valX, valY, valZ, rrlA, rrlC-rrlH, rrsA, rrsC-rrsH, rrfA*, and *rrfC-rrfH*) using the Bedtools Maskfasta option. Previously we have identified regions of significant enrichment after isoleucine starvation by False Discovery Rate (FDR) analysis using pyCalculateFDR.py^2^. Selected regions including the A-site finger (ASF, nucleotide 834-927 in 23S rRNA), Sarcin-Ricin Loop (SRL, nucleotide 2652-2673 in 23S rRNA), Helix 15 (H15, nucleotide 328-407 in 16S rRNA) and tRNA^IleTUV^ were used to calculate the normalized cDNA coverage. Deletions and substitutions introduced in the cDNA reads were counted in selected regions showing significant enrichment after isoleucine starvation. cDNA reads and crosslinking sites were visualized by plots, boxplots and heat maps in R.

### Data availability

Sequencing data has been deposited with GEO accession GSE150416 Samples RelA_0_Ex1, RelA_30_Ex1, RelA_0_Ex2 and RelA_30_Ex2 are from ^2^ and can be accessed with GEO accession: GSM2912989, GSM2912991, GSM2912990 and GSM2912992.

## Acknowledgements

We thank Kenn Gerdes and Sine Lo Svenningsen for critical reading of the manuscript and in providing important suggestions.

## Author contributions

K.S.W conceived and initiated the study. K.S.W and A.K.S. performed the experiments and analyzed the data. K.S.W and A.K.S. wrote the paper.

