## Supplementary information for "The RelA hydrolase domain: a molecular switch for (p)ppGpp synthesis"

**Strains Constructed**

Oligos, strains and plasmids used for strain construction are shown in Supplementary Table S1

**MG1655*relA*$^{QUAD}$ and MG1655*relA*$^{QUAD}$*::HTF***

A chloramphenicol cassette was inserted into isoleucine 116 of *relA* in MG1655 and MG1655 *relA::HTF*. The cassette was generated by PCR using plasmid pWRG100 as template and oligos I116-CM-f and I116-CM-rv and was electroporated (app. 200ng) into MG1655 and MG1655 relA::HTF expressing lambda red recombinase from plasmid pWRG99 [1]. After 1 hr of phenotypic expression cells and were plated onto LB agar plates containing 10µg/mL chloramphenicol and 100µg/mL ampicillin and grown at 30ºC. Insertion of cassette generated MG1655 *relA*$^{I116::cm}$ and MG1655 *relA*$^{I116::cm}$*::HTF,* which were confirmed by PCR. Alanine substitutions in R117, Q118, K120 and H123 (referred to as QUAD) were introduced by replacing the cassette in MG1655 *relA*$^{I116::cm}$ and MG1655 *relA*$^{I116::CM}$*::HTF* with a double stranded DNA oligonucleotides containing substitution mutations. Double stranded DNA was generated by hybridization of 500 pmol of quad-scarless-f and quad-scarless-rv oligonucleotides in a reaction containing 10mM Tris-HCl pH 7.5, 100mM NaCl and 1mM EDTA followed by a subsequent desalting step using a G25 Spin column (GE healthcare). 25 pmol of double stranded DNA oligonucleotide was electroporated into MG1655 *relA*$^{I116::cm}$ and MG1655 *relA*$^{I116::cm}$*::HTF* expressing lambda recombinase from pWRG99 and after 1hr of phenotypic expression serially diluted and plated on LB-plates containing 100µg/mL ampicillin and 1µg/mL anhydrotetracycline. Positive Sce-I resistant clones were re-streaked and sequenced.

**MG1655 *relA*$^{\Delta 116\text{-}129}$ and MG1655 *relA*$^{\Delta 116\text{-}129}$*::HTF***

Deletions of residues 116-129 in relA (referred to as Δ116-129) were constructed as above using hybridized oligos Delta116-129-scarless-f and Delta116-129-scarless-rv.

**MG1655 *relA*$^{I116L}$ and MG1655 *relA*$^{I116L}$*::HTF***

Isoleucine in position 116 was substituted for Leucine (referred to as I116L) as above using hybridized oligos I116L-scarless-f and I116L-scarless-rv.

**MG1655 *relA*$^{A121E}$ and MG1655 *relA*$^{A121E}$*::HTF***

Alanine in position 121 was substituted for glutamate (referred to as A121E) as described above using oligos A121E-scarless-f and A121E-scarless-rv.

**MG1655 *relA*$^{L119M}$ and MG1655 *relA*$^{L119M}$*::HTF***

Leucine 119 of RelA was substituted for methionine (referred to as L119M) as described above using hybridized oligos L119M-scarless-f and L119M-scarless-rv

**MG1655 *relA*$^{M113K}$ and MG1655 *relA*$^{M113K}$*::HTF***

Methionine 113 was substituted for lysine (referred to as M113K) as described above using hybridized oligos M113K-scarless-f and M113K-scarless-rv

**MG1655 *relA*$^{Q118L}$ and MG1655 *relA*$^{Q118L}$*::HTF***

Glutamine 118 was substituted for leucine (referred to as Q118L) as described above using oligos Q118L-scarless-f and Q118L-scarless-rv.

**MG1655 *relA$^{H108AR111A}$::HTF***

Histidine 108 and arginine 111 was substituted for alanine (referred to as H108AR111A) in MG1655*relA::HTF* as described above using oligos H108AR111A-scarless-f and H108AR111A –scarless-rv.

**MG1655 *relA$^{W39A}$::HTF***

A chloramphenicol cassette was inserted into tryptophan W39 of relA in MG1655*relA::HTF*. The cassette was generated by PCR using plasmid pWRG100 as template and oligos W39-CM-f and W39-CM-rv and was electroporated MG1655*relA::HTF* as described previously. W39 was substituted for alanine (referred to as W39A) by replacing the cassette in MG1655*relA$^{W39::cm}$::HTF* as described above using oligos W39A-scarless-f and W39A-scarless-rv.

**MG1655 *relA$^{\Delta W39}$::HTF***

Deletion of W39 (referred to as ΔW39) was introduced as described above using oligos DeltaW39-scarless-f and DeltaW39-scarless-rv.

**MG1655 *relA$^{R96AK101A}$::HTF***

A chloramphenicol cassette was inserted into serine S98 of relA in MG1655*relA::HTF* as described above using a PCR product generated from oligos S98-CM-f and S98-CM-rv. R96 and K101 were substituted for alanine (referred to as R96AK101A) by replacing the cassette in MG1655*relA$^{S98::cm}$::HTF* with hybridized oligos R96AK101A-scarless-f and R96AK101A-scarless-rv as described previously.

**MG1655 *relA$^{R136AR137A}$::HTF***

A chloramphenicol cassette was inserted into arginine R136 of relA in MG1655*relA::HTF* as described previously using a PCR product generated by oligos R136-CM-f and R136-CM-rv R136 and R137 were substituted for alanine (referred to as R136AR137A) by replacing the cassette in MG1655*relA$^{R136::cm}$::HTF* with a hybridized oligos R136AR137A-scarless-f and R136AR137A-scarless-rv as described above.

**Protein stability by Western blotting analysis**

Cultures of MG1655 *ΔrelA*, MG1655 *relA::HTF*, MG1655 *relA$^{A121E}$::HTF*, MG1655 *relA$^{QUAD}$::HTF*, MG1655 *relA$^{I116L}$::HTF* and MG1655 *relA$^{Δ116-129}$::HTF* were grown exponentially in MOPS minimal medium supplemented with 0.2% glucose and all nucleobases (10 μg $ml^{-1}$ of each) at 30ºC. At $OD_{600}$ = 0.2 Isoleucine starvation was induced by addition of L-Valine to a final concentration of 500μg/mL. A 1mL sample was collected before and after 30 min of Isoleucine starvation and the cells pelleted by centrifugation at 4ºC. The pellet was resuspend in 50μL 1x LDS loading buffer (Life technologies) with 5mM DTT and boiled for 5 min. After a brief step of centrifugation at 14krpm, 20μL of sample was loaded onto a 4-12% NuPAGE gel (Life technologies) and the protein separated by electrophoresis in 1xMOPS running buffer (Life technologies). The protein was transferred to a PVDF membrane (Amersham) and six Histidine tagged RelA protein detected by incubation with pentahis primary antibodies (Qiagen) followed by incubation with HRP conjugated mouse IgG secondary antibodies (Sigma) in PBS with 0.1% Tween-20 and 5% Milk powder. The protein bands were visualized using Pierce ECL chemiluminiscense substrate (Thermo scientific) according to manufacturer's instructions and the signal detected in an Imagequant LAS4100.

### Supplementary Table S1

| Oligos | |
|---|---|
| I116-CM-f | GGTCGTTAACCTTATTCACGGCGTGCGTGATATGGCGGCGCTAGACTATATTACCCTGTT |
| I116-CM-rv | CGGAGGAAACAGAATCAGTGTGCGTCGCTTTCAGCTGGCGCGCCTTACGCCCCGCCCTGC |
| Quad_scarless-f | TAACCTTATTCACGGCGTGCGTGATATGGCGGCGATCGCGGCCCTGGCGGCGACGGCCACTGATTCTGTTTCCTCCGAACAGGTCGATAACG |
| Quad_scarless-rv | CGTTATCGACCTGTTCGGAGGAAACAGAATCAGTGGCCGTCGCCGCCAGGGCCGCGATCGCCGCCATATCACGCACGCCGTGAATAAGGTTA |
| Delta116-129_scarless-f | GGTCGTTAACCTTATTCACGGCGTGCGTGATATGGCGGCGGAACAGGTCGATAACGTTCGCCGGATGTTATTGGCGATGG |
| Delta116-129_scarless-rv | CCATCGCCAATAACATCCGGCGAACGTTATCGACCTGTTCCGCCGCCATATCACGCACGCCGTGAATAAGGTTAACGACC |
| I116L-scarless-f | GTTAACCTTATTCACGGCGTGCGTGATATGGCGGCGCTGCGCCAGCTGAAAGCGACGCACACTGATTCTGTTTCCTCCGA |
| I116L-scarless-rv | TCGGAGGAAACAGAATCAGTGTGCGTCGCTTTCAGCTGGCGCAGCGCCGCCATATCACGCACGCCGTGAATAAGGTTAAC |
| A121E-scarless-f | TTAACCTTATTCACGGCGTGCGTGATATGGCGGCGATCCGCCAGCTGAAAGAAACGCACACTGATTCTGTTTCCTCCGAACAGGTCGA |
| A121E-scarless-rv | TCGACCTGTTCGGAGGAAACAGAATCAGTGTGCGTTTCTTTCAGCTGGCGGATCGCCGCCATATCACGCACGCCGTGAATAAGGTTAA |
| L119M- | TTAACCTTATTCACGGCGTGCGTGATATGGCGGCGATCCGCCAGATGAAAGCGACGCACAC |

| | |
|---|---|
| scarless-f | TGATTCTGTTTCCTCCGAACA |
| L119M-scarless-rv | TGTTCGGAGGAAACAGAATCAGTGTGCGTCGCTTTCATCTGGCGGATCGCCGCCATATCACGCACGCCGTGAATAAGGTTAA |
| M113K-scarless-f | GTCGGTCGTTAACCTTATTCACGGCGTGCGTGATAAAGCGGCGATCCGCCAGCTGAAAGCGACGCACACTGATTCTGTTTC |
| M113K-scarless-rv | GAAACAGAATCAGTGTGCGTCGCTTTCAGCTGGCGGATCGCCGCTTTATCACGCACGCCGTGAATAAGGTTAACGACCGAC |
| Q118L-scarless-f | AACCTTATTCACGGCGTGCGTGATATGGCGGCGATCCGCCTGCTGAAAGCGACGCACACTGATTCTGTTTCCTCCG |
| Q118L-scarless-rv | CGGAGGAAACAGAATCAGTGTGCGTCGCTTTCAGCAGGCGGATCGCCGCCATATCACGCACGCCGTGAATAAGGTT |
| H108AR111A-scarless-f | GTGAGAGCGTCGGTAAGTCGGTCGTTAACCTTATTGCGGGCGTGGCGGATATGGCGGCGATCCGCCAGCTGAAAGCGACGCACACTGATTCTG |
| H108AR111A-scarless-rv | CAGAATCAGTGTGCGTCGCTTTCAGCTGGCGGATCGCCGCCATATCCGCCACGCCCGCAATAAGGTTAACGACCGACTTACCGACGCTCTCAC |
| W39-CM-f | CCAGCCAGAAGTCGTGTGAGTGCTTAGCCGAAACCCTAGACTATATTACCCTGTT |
| W39-CM-rv | CATCCGGATGCCCCTGCGTCTGTTGCAGACAATACGCCGCCTTACGCCCCGCCCTGC |
| W39A-scarless-f | GTATTACCAGCCAGAAGTCGTGTGAGTGCTTAGCCGAAACCGCCGCGTATTGTCTGCAACAGACGCAGGGGCATCCGGATG |
| W39A-scarless-rv | CATCCGGATGCCCCTGCGTCTGTTGCAGACAATACGCGGCGGTTTCGGCTAAGCACTCACACGACTTCTGGCTGGTAATAC |
| DeltaW39-scarless-f | CCAGCCAGAAGTCGTGTGAGTGCTTAGCCGAAACCGCGTATTGTCTGCAACAGACGCAGGGGCATCCGGA |
| DeltaW39-scarless-f | TCCGGATGCCCCTGCGTCTGTTGCAGACAATACGCGGTTTCGGCTAAGCACTCACACGACTTCTGGCTGG |

| | |
|---|---|
| S98-CM-f | GATGCCAACGTAGTCAGCGAAGATGTGCTGCGTGAGCTAGACTATATTACCCTGTT |
| S98-CM-rv | CACGCCGTGAATAAGGTTAACGACCGACTTACCGACCGCCTTACGCCCCGCCCTGC |
| R96AK101A-scarless-f | CTGGCGGATGCCAACGTAGTCAGCGAAGATGTGCTGGCGGAGAGCGTCGGTGCGTCGGTCGTTAACCTTATTCACGGCGTGCGTGATATG |
| R96AK101A-scarless-rv | CATATCACGCACGCCGTGAATAAGGTTAACGACCGACGCACCGACGCTCTCCGCCAGCACATCTTCGCTGACTACGTTGGCATCCGCCAG |
| R136-CM-f | CTGATTCTGTTTCCTCCGAACAGGTCGATAACGTTCTAGACTATATTACCCTGTT |
| R136-CM-rv | CAGCGAAAATCATCGACCATCGCCAATAACATCCGCGCCTTACGCCCCGCCCTGC |
| R136AR137A-scarless-f | CTGATTCTGTTTCCTCCGAACAGGTCGATAACGTTGCGGCCATGTTATTGGCGATGGTCGATGATTTTCGCTGCG |
| R136AR137A-scarless-rv | CGCAGCGAAAATCATCGACCATCGCCAATAACATGGCCGCAACGTTATCGACCTGTTCGGAGGAAACAGAATCAG |
| Loop-mut-f | CGGTCGTTAACCTTATTCAC |
| Loop-mut-rv | CTACGCAGCGAAAATCATC |
| miRCat-33 | rAppTGGAATTCTCGGGTGCCAAGG/ddC/ |
| 33-rev | CCTTGGCACCCGAGAATT |
| L5Aa | InvddT/**ACAC**rGrArCrGrCrUrCrUrUrCrCrGrArUrCrUrNrNrNrUrArArGrC |
| L5Ab | InvddT/**ACAC**rGrArCrGrCrUrCrUrUrCrCrGrArUrCrUrNrNrNrArUrUrArGrC |
| L5Ad | InvddT/**ACAC**rGrArCrGrCrUrCrUrUrCrCrGrArUrCrUrNrNrNrCrGrCrUrUrArGrC |
| L5Bb | InvddT/**ACAC**rGrArCrGrCrUrCrUrUrCrCrGrArUrCrUrNrNrNrGrUrGrArGrC |
| L5Bc | InvddT/**ACAC**rGrArCrGrCrUrCrUrUrCrCrGrArUrCrUrNrNrNrCrArCrUrArGrC |
| L5Bd | InvddT/**ACAC**rGrArCrGrCrUrCrUrUrCrCrGrArUrCrUrNrNrNrUrCrUrCrUrArGrC |
| P5 | AATGATACGGCGACCACCGAGATCTACACTCTTTCCCTACACGACGCTCTTCCGATCT |
| PE | CAAGCAGAAGACGGCATACGAGATCGGTCTCGGCATTCCTGGCCTTGGCACCCGAGAATTCC |
| Strains | |

| | |
|---|---|
| MG1655 *relA::HTF* | [2] |
| MG1655 *ΔrelA* | [3] |
| Plasmids | |
| pWRG99 | [1] |
| pWRG100 | [1] |

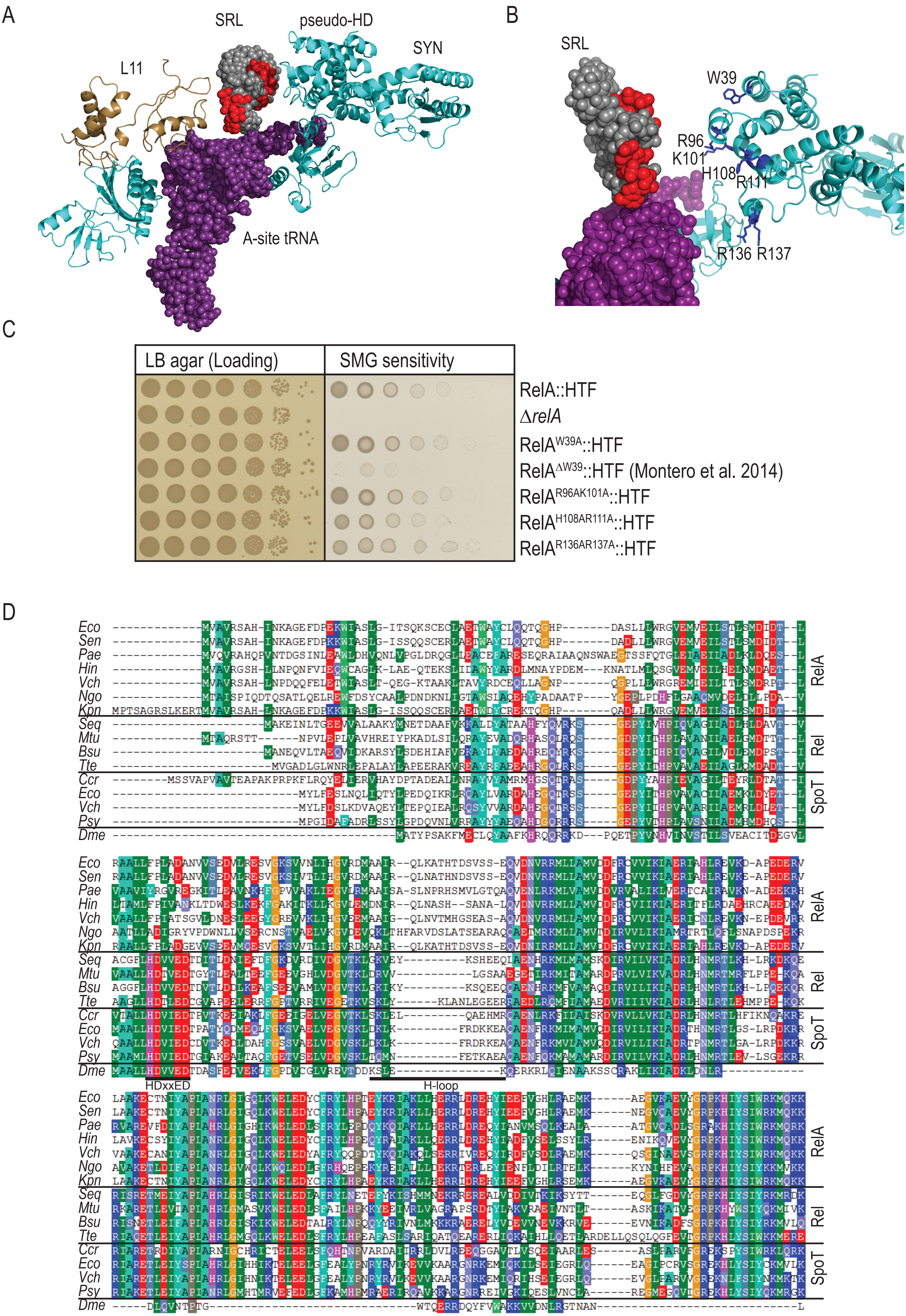
A
SRL
pseudo-HD
SYN
L11
A-site tRNA
B
SRL
W39
R96
K101
H108
R111
R136 R137
C
LB agar (Loading)
SMG sensitivity
RelA::HTF
ΔrelA
RelA^W39A::HTF
RelA^ΔW39::HTF (Montero et al. 2014)
RelA^R96AK101A::HTF
RelA^H108AR111A::HTF
RelA^R136AR137A::HTF
D
Eco
Sen
Pae
Hin
Vch
Ngo
Kpn
Seq
Mtu
Bsu
Tte
Ccr
Eco
Vch
Psy
Dme
RelA
Rel
SpoT
HDxxED
H-loop

Fig.S1 - Page 1

E

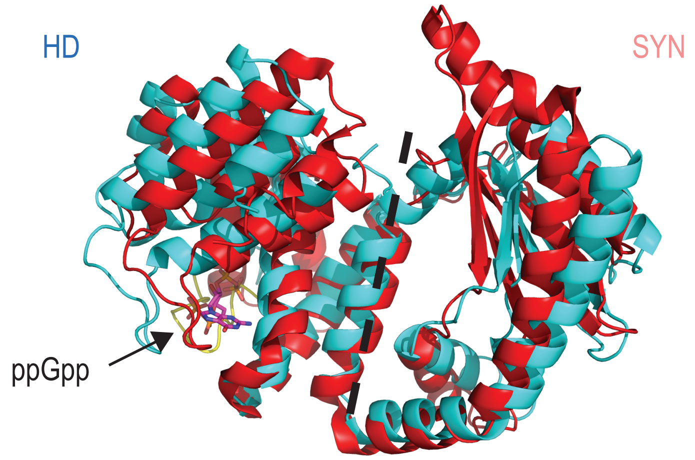

F

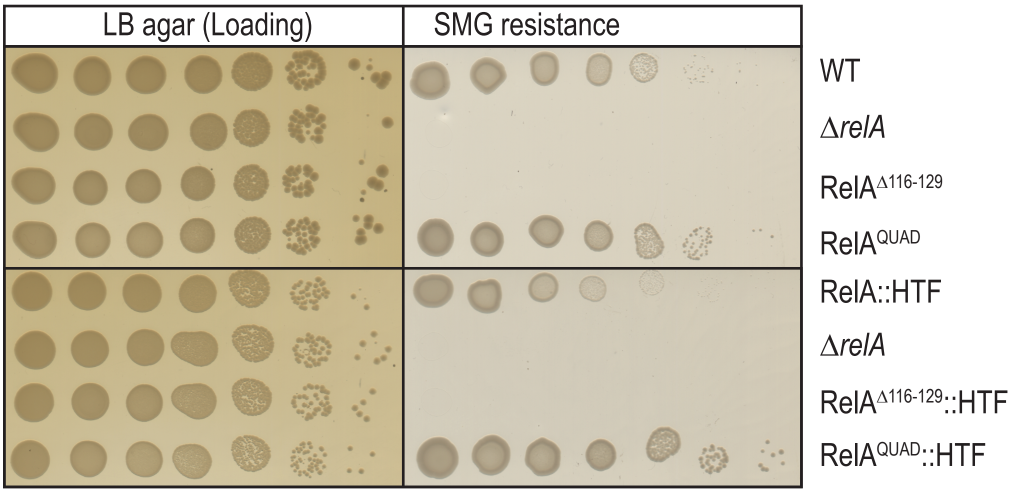

G

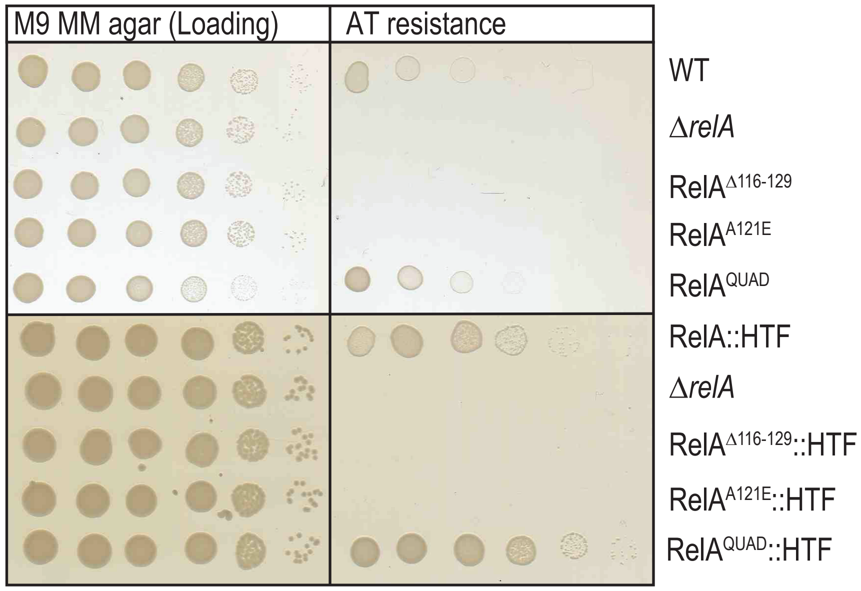

H

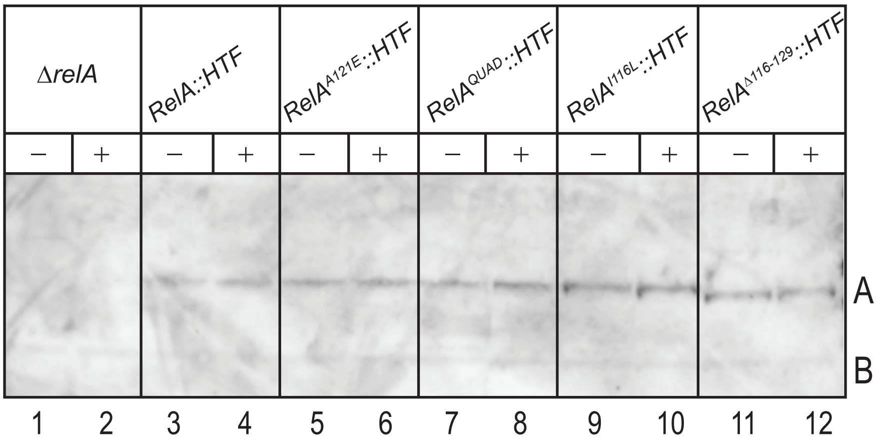

Fig. S1 - Page 2

**Supplementary Fig. S1** Identification of the H-loop in the hydrolase domain. **A**) Structure of RelA (cyan) bound with uncharged tRNA (magenta) at the ribosomal A-site (PDB: 5IQR). RelA forms an elongated structure on the ribosome enclosing the A-site tRNA. Potential RelA interaction sites with the Sarcin-Ricin Loop (SRL) are indicated in red and ribosomal protein L11 is shown in brown. **B**) Close-up on residues in RelA pseudo-HD (blue) predicted to interact with the Sarcin-Ricin Loop (SRL). **C**) Assaying functionality of RelA hydrolase substitution mutants. MG1655 *relA::HTF*, *ΔrelA*, $relA^{W39A}$*::HTF*, $relA^{\Delta W39}$*::HTF*, $relA^{R96AK101A}$*::HTF*, $relA^{H108AR111A}$*::HTF* and $relA^{R136AR137A}$*::HTF* were grown in LB medium. The cells were washed in PBS, serial diluted and spotted on loading control plates (LB agar) and MOPS MM SMG plates (SMG resistance) and grown at 30ºC. **D**) Multiple sequence alignment of selected RelA, Rel and SpoT N-terminal sequences. *Eco, Escherichia coli, Sen, Salmonella enterica, Pae, Pseudomonas aeruginosa, Hin, Hemophilus influenza, Vch, Vibrio cholera, Ngo, Neisseria gonorrhoeae, Kpn, Klebsiella pneumoniae, Seq, Streptococcus dysgalactiae subsp. equisimilis, Mtu, Mycobacterium tuberculosis, Bsu, Bacillus subtilis,Tte, Thermus thermophilus, Ccr, Caulobacter crescentus, Psy, Pseudomonas syringae, Dme, Drosophila melanogaster*. Location of the H-loop is indicated with a box and conserved metal-dependent pyrophosphohydrolase HDXXED motif is underlined. **E**) Overlay of N-terminal domain (hydrolase and synthetase) of $Rel_{Tte}$ from *Thermus thermophilus*(PDB: 6S2T shown in red) with N-terminal domain of RelA (PDB: 5IQR). Location of ppGpp in $Rel_{Tte}$ structure are indicated. The H-loop in RelA is indicated in yellow and the dotted line separates the two functional domains. **F**) Functionality test of tagged and untagged RelA H-loop deletion mutants. MG1655 (WT), MG1655 *ΔrelA*, MG1655 $relA^{\Delta 116\text{-}129}$ and $relA^{QUAD}$ or HTF-tagged (Cterminal six histidine, TEV cleavage site and three FLAG epitopes) versions were grown overnight in LB medium at 37ºC. The cultures were then washed in PBS serial diluted and plated onto LB agar (loading) and MOPS MM SMG plates (SMG resistance). Un-tagged strains were grown at 37ºC and tagged strains at 30ºC. **G)** MG1655 (WT), MG1655 *ΔrelA*, MG1655 *relA*$^{\Delta 116\text{-}129}$, MG1655 *relA*$^{A121E}$ and *relA*$^{QUAD}$ or HTF-tagged versions were grown and diluted as in F) and spotted onto M9 minimal agar plates containing all amino acids except histidine (FN19), 1mM adenine, 1mM thiamine without or with 15mM 3-amino-1,2,3-triazole (AT resistance). Un-tagged strains were grown at 37ºC and tagged strains at 30ºC. **H**) Detection of chromosomally encoded HTF-tagged RelA. MG1655 ΔrelA (Control), MG1655 *relA::HTF*, *relA*$^{A121E}$*::HTF*, *relA*$^{QUAD}$*::HTF*, *relA*$^{I116L}$*::HTF* and *relA*$^{\Delta 116\text{-}129}$*::HTF* were grown exponentially in MOPS minimal medium at 30ºC. Samples were collected before and 30 min isoleucine starvation. HTF tagged protein was detected by western analysis using penta-his antibodies as described in methods. A indicates position of RelA::HTF, B indicates position of unspecific band present in all samples, which can be used as a loading control.

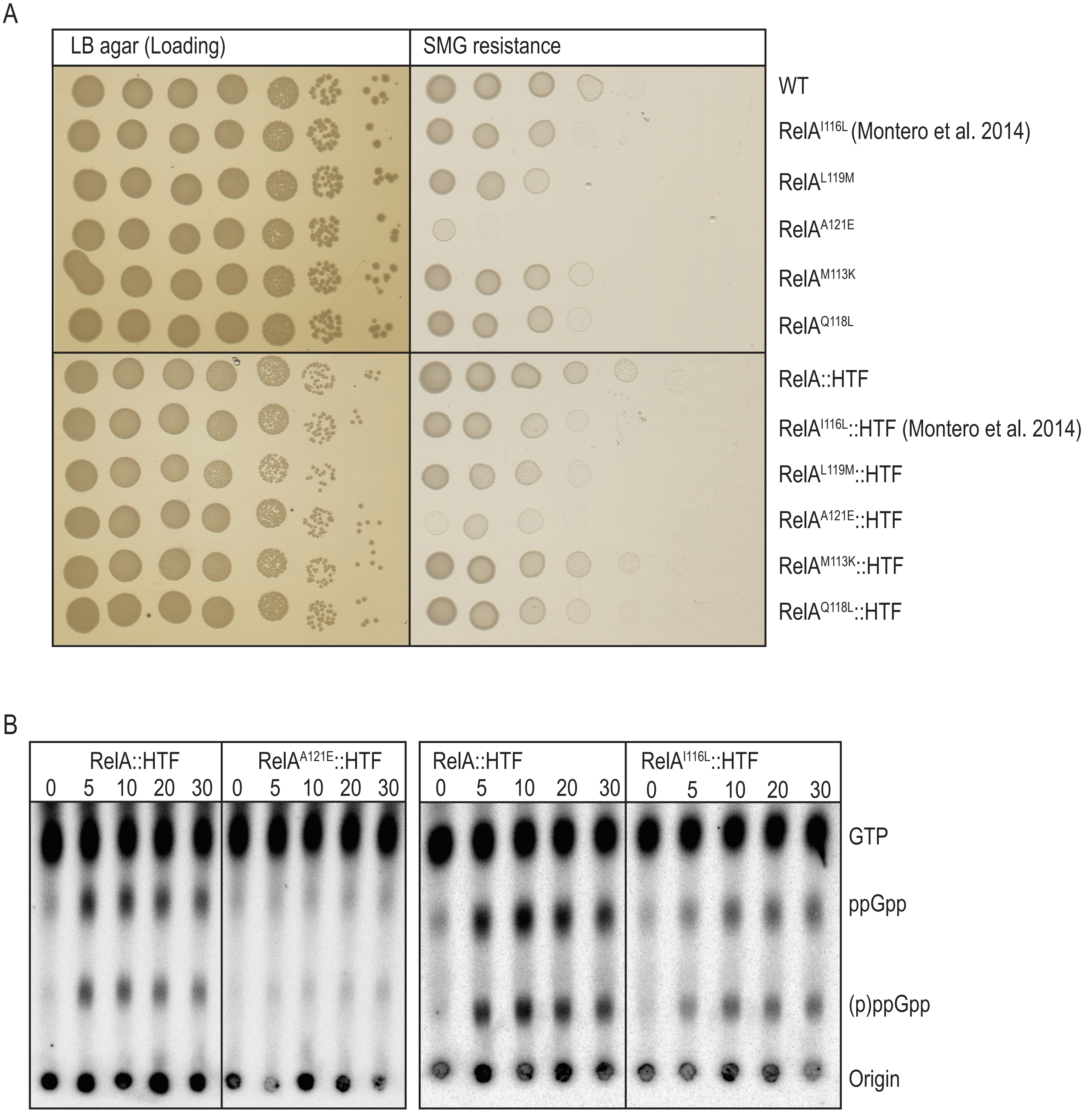
A
LB agar (Loading)
SMG resistance
WT
RelA^I116L (Montero et al. 2014)
RelA^L119M
RelA^A121E
RelA^M113K
RelA^Q118L
RelA::HTF
RelA^I116L::HTF (Montero et al. 2014)
RelA^L119M::HTF
RelA^A121E::HTF
RelA^M113K::HTF
RelA^Q118L::HTF
B
RelA::HTF
RelA^A121E::HTF
RelA::HTF
RelA^I116L::HTF
0 5 10 20 30
GTP
ppGpp
(p)ppGpp
Origin

Fig. S2

**Supplementary Fig. S2** Substitution mutations in the H-loop modulates (p)ppGpp synthesis. **A)** Functionality testing of untagged or tagged RelA H-loop substitution mutants. MG1655 (WT), *ΔrelA*, *relA*$^{I116L}$, *relA*$^{L119M}$, *relA*$^{A121E}$, *relA*$^{M113K}$ and *relA*$^{Q118L}$ or HTF-tagged versions were grown overnight in LB medium at 37ºC. The cultures were then washed in PBS serial diluted and plated onto LB agar (Loading) and MOPS MM SMG plates (SMG resistance). Un-tagged strains were grown at 37ºC and tagged strains at 30ºC. **B)** (p)ppGpp measurements of MG1655 *relA::HTF, relA*$^{A121E}$*::HTF* and *relA*$^{I116L}$*::HTF*. Cells were grown exponentially in MOPS minimal medium containing $^{32}$P-labeled phosphate at 30ºC. Isoleucine starvation was induced by addition of L-valine, to a final concentration of 500µg/mL. Samples were collected before (time zero) and after starvation, precipitated and separated by thin layer chromatography. Position of GTP, ppGpp and pppGpp is indicated.

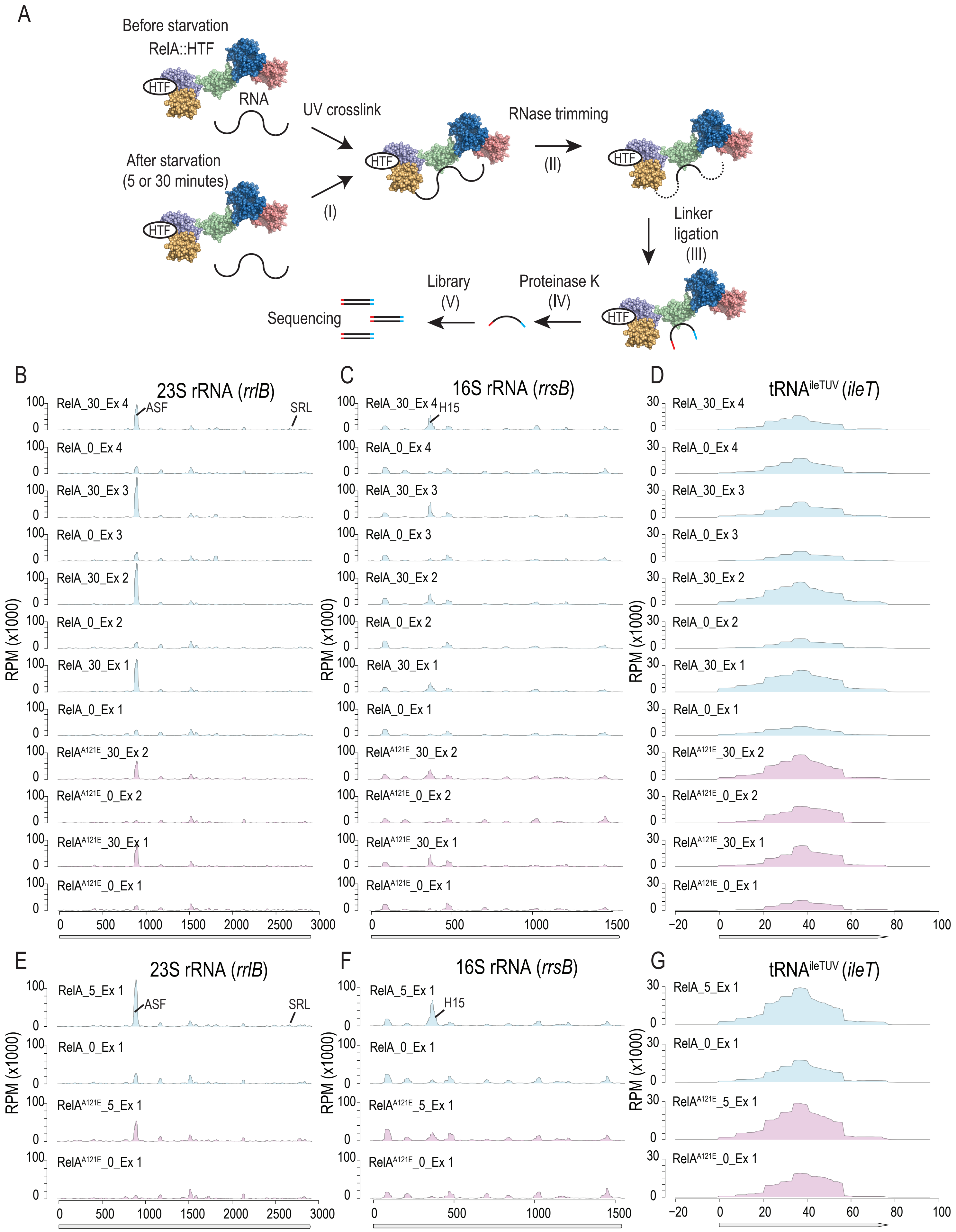
A
Before starvation
RelA::HTF
HTF
RNA
After starvation
(5 or 30 minutes)
UV crosslink
(I)
RNase trimming
(II)
Linker ligation
(III)
Proteinase K
(IV)
Library
(V)
Sequencing
B
23S rRNA (rrlB)
ASF
SRL
RPM (x1000)
RelA_30_Ex 4
RelA_0_Ex 4
RelA_30_Ex 3
RelA_0_Ex 3
RelA_30_Ex 2
RelA_0_Ex 2
RelA_30_Ex 1
RelA_0_Ex 1
RelA^A121E_30_Ex 2
RelA^A121E_0_Ex 2
RelA^A121E_30_Ex 1
RelA^A121E_0_Ex 1
C
16S rRNA (rrsB)
H15
D
tRNA^ileTUV (ileT)
E
23S rRNA (rrlB)
RelA_5_Ex 1
RelA_0_Ex 1
RelA^A121E_5_Ex 1
RelA^A121E_0_Ex 1
F
16S rRNA (rrsB)
G
tRNA^ileTUV (ileT)

Fig. S3

**Supplementary Fig. S3** RelA-RNA interactions by Crosslinking and analysis of cDNAs (CRAC). **A)** Method overview. Step (I): UV-irradiation of cell samples before and 5 or 30 minutes after amino acid starvation. Step (II): Purification of RelA-RNA complexes via the FLAG epitope tag and RNA trimming by RNases. Step (III): Immobilization of complexes via the His-tag and DNA linker ligation to RNA 3'- and 5'-ends. Step (IV): Size selection of RelA-RNA complexes and RelA degradation by protease K treatment. Step (V): Finally, the cDNA libraries were generated by RT-PCR and subjected to deep sequencing. **B**) DNA reads mapping to 23S rRNA (*rrlB*) obtained from CRAC analysis of MG1655 *relA::HTF and relA*$^{A121E}$*::HTF* before (0) and after isoleucine starvation (number indicates time in minutes). Reads are normalized to Reads per million (RPM). **C)** Reads aligning to 16S rRNA (*rrsB*). **D)** Reads aligning to isoleucine tRNA (*ileT*). **E**) Normalized Reads obtained after short-term starvation (5 min) aligned to 23S rRNA **F**) 16S rRNA or **G**) isoleucine tRNA. Position of A-site finger (ASF), Sarcin-ricin Loop (SRL) and Helix 15 (H15) are indicated with arrows. Each experiment (Ex) refers to an independent biological replicate.

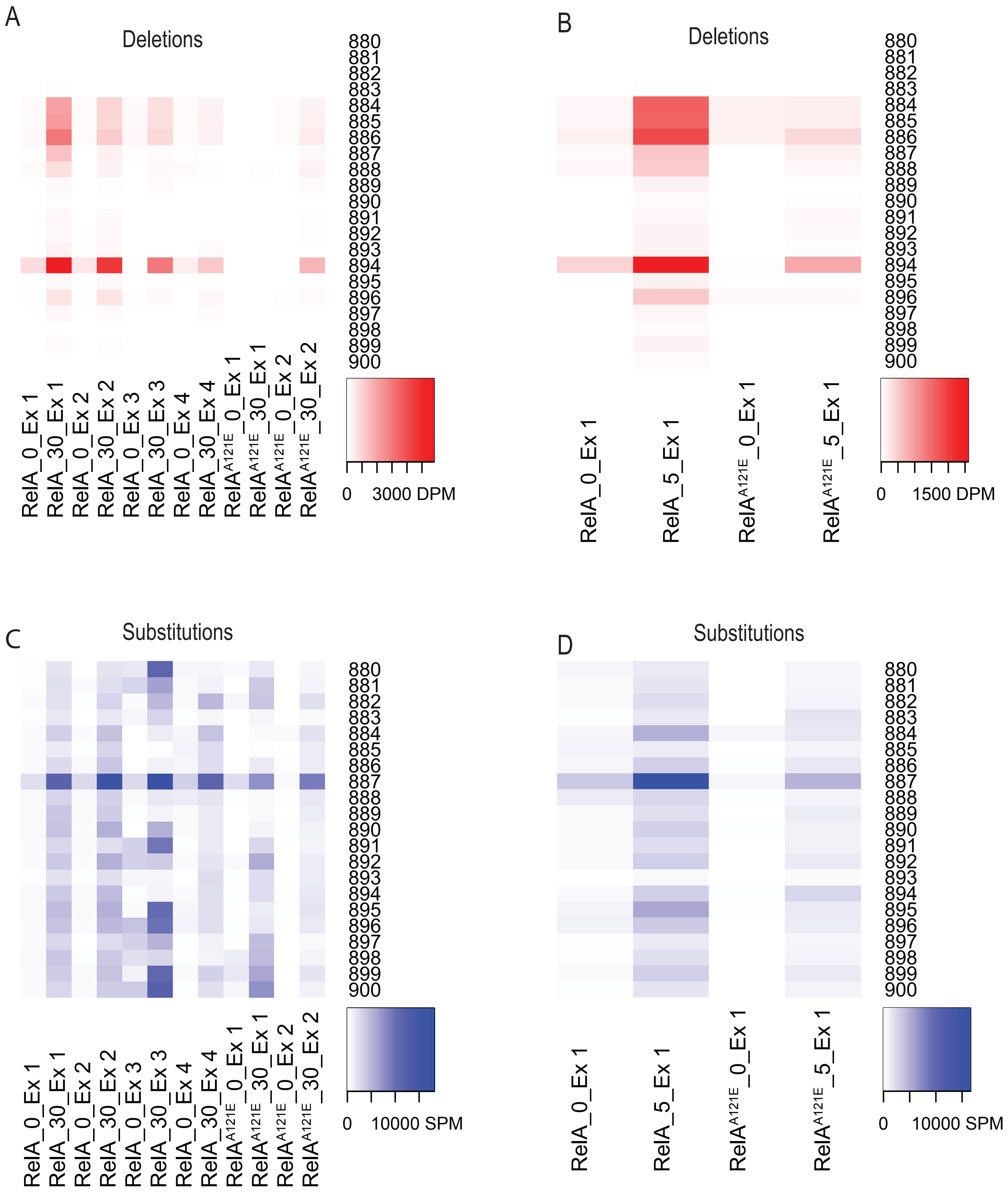
A
Deletions
RelA_0_Ex 1
RelA_30_Ex 1
RelA_0_Ex 2
RelA_30_Ex 2
RelA_0_Ex 3
RelA_30_Ex 3
RelA_0_Ex 4
RelA_30_Ex 4
RelA$^{A121E}$_0_Ex 1
RelA$^{A121E}$_30_Ex 1
RelA$^{A121E}$_0_Ex 2
RelA$^{A121E}$_30_Ex 2
0
3000 DPM
B
Deletions
RelA_0_Ex 1
RelA_5_Ex 1
RelA$^{A121E}$_0_Ex 1
RelA$^{A121E}$_5_Ex 1
0
1500 DPM
C
Substitutions
RelA_0_Ex 1
RelA_30_Ex 1
RelA_0_Ex 2
RelA_30_Ex 2
RelA_0_Ex 3
RelA_30_Ex 3
RelA_0_Ex 4
RelA_30_Ex 4
RelA$^{A121E}$_0_Ex 1
RelA$^{A121E}$_30_Ex 1
RelA$^{A121E}$_0_Ex 2
RelA$^{A121E}$_30_Ex 2
0
10000 SPM
D
Substitutions
RelA_0_Ex 1
RelA_5_Ex 1
RelA$^{A121E}$_0_Ex 1
RelA$^{A121E}$_5_Ex 1
0
10000 SPM

Fig. S4 - Page 1

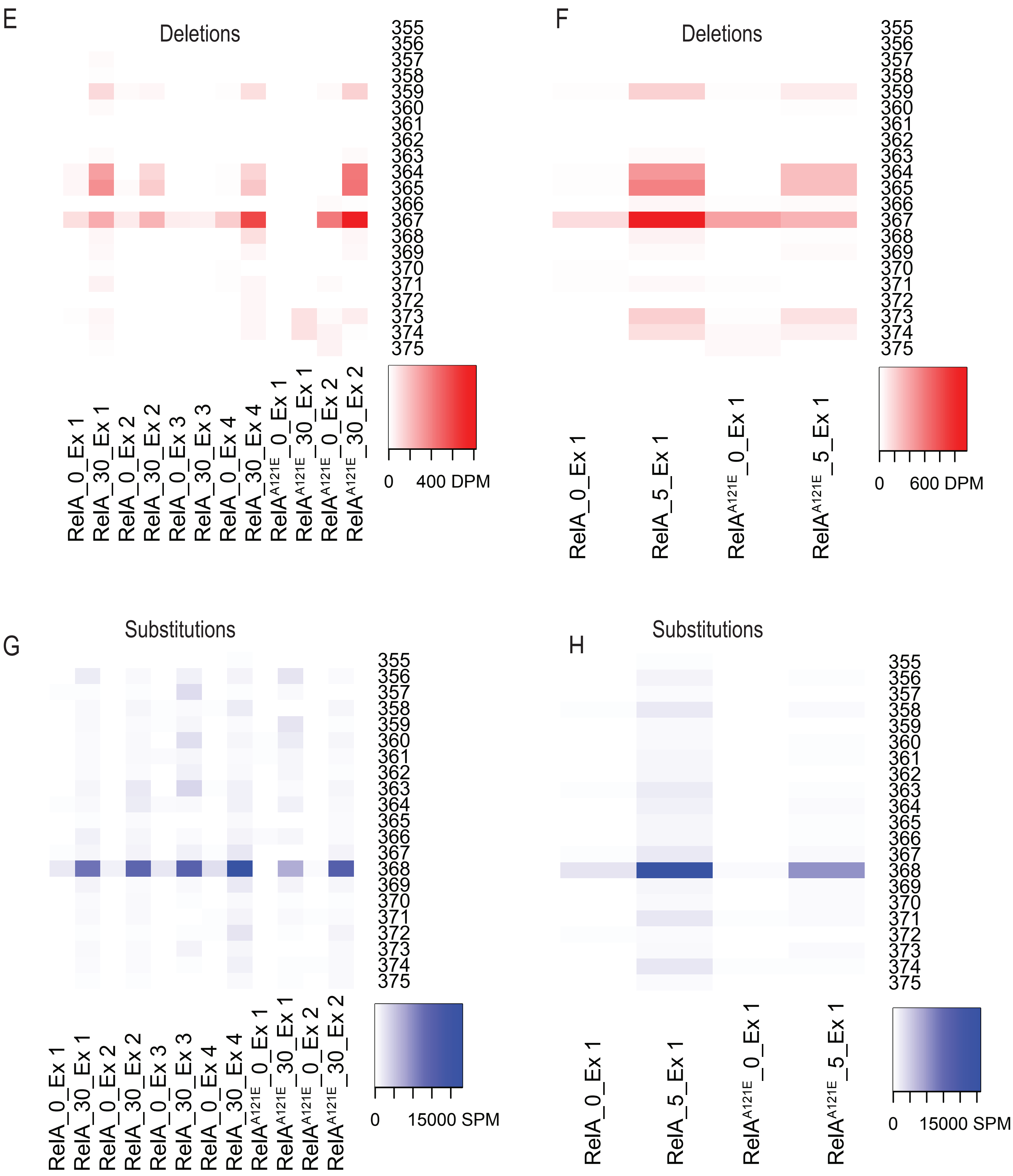
E
Deletions
355 356 357 358 359 360 361 362 363 364 365 366 367 368 369 370 371 372 373 374 375
RelA_0_Ex 1
RelA_30_Ex 1
RelA_0_Ex 2
RelA_30_Ex 2
RelA_0_Ex 3
RelA_30_Ex 3
RelA_0_Ex 4
RelA_30_Ex 4
RelA$^{A121E}$_0_Ex 1
RelA$^{A121E}$_30_Ex 1
RelA$^{A121E}$_0_Ex 2
RelA$^{A121E}$_30_Ex 2
0 400 DPM
F
Deletions
355 356 357 358 359 360 361 362 363 364 365 366 367 368 369 370 371 372 373 374 375
RelA_0_Ex 1
RelA_5_Ex 1
RelA$^{A121E}$_0_Ex 1
RelA$^{A121E}$_5_Ex 1
0 600 DPM
G
Substitutions
355 356 357 358 359 360 361 362 363 364 365 366 367 368 369 370 371 372 373 374 375
RelA_0_Ex 1
RelA_30_Ex 1
RelA_0_Ex 2
RelA_30_Ex 2
RelA_0_Ex 3
RelA_30_Ex 3
RelA_0_Ex 4
RelA_30_Ex 4
RelA$^{A121E}$_0_Ex 1
RelA$^{A121E}$_30_Ex 1
RelA$^{A121E}$_0_Ex 2
RelA$^{A121E}$_30_Ex 2
0 15000 SPM
H
Substitutions
355 356 357 358 359 360 361 362 363 364 365 366 367 368 369 370 371 372 373 374 375
RelA_0_Ex 1
RelA_5_Ex 1
RelA$^{A121E}$_0_Ex 1
RelA$^{A121E}$_5_Ex 1
0 15000 SPM

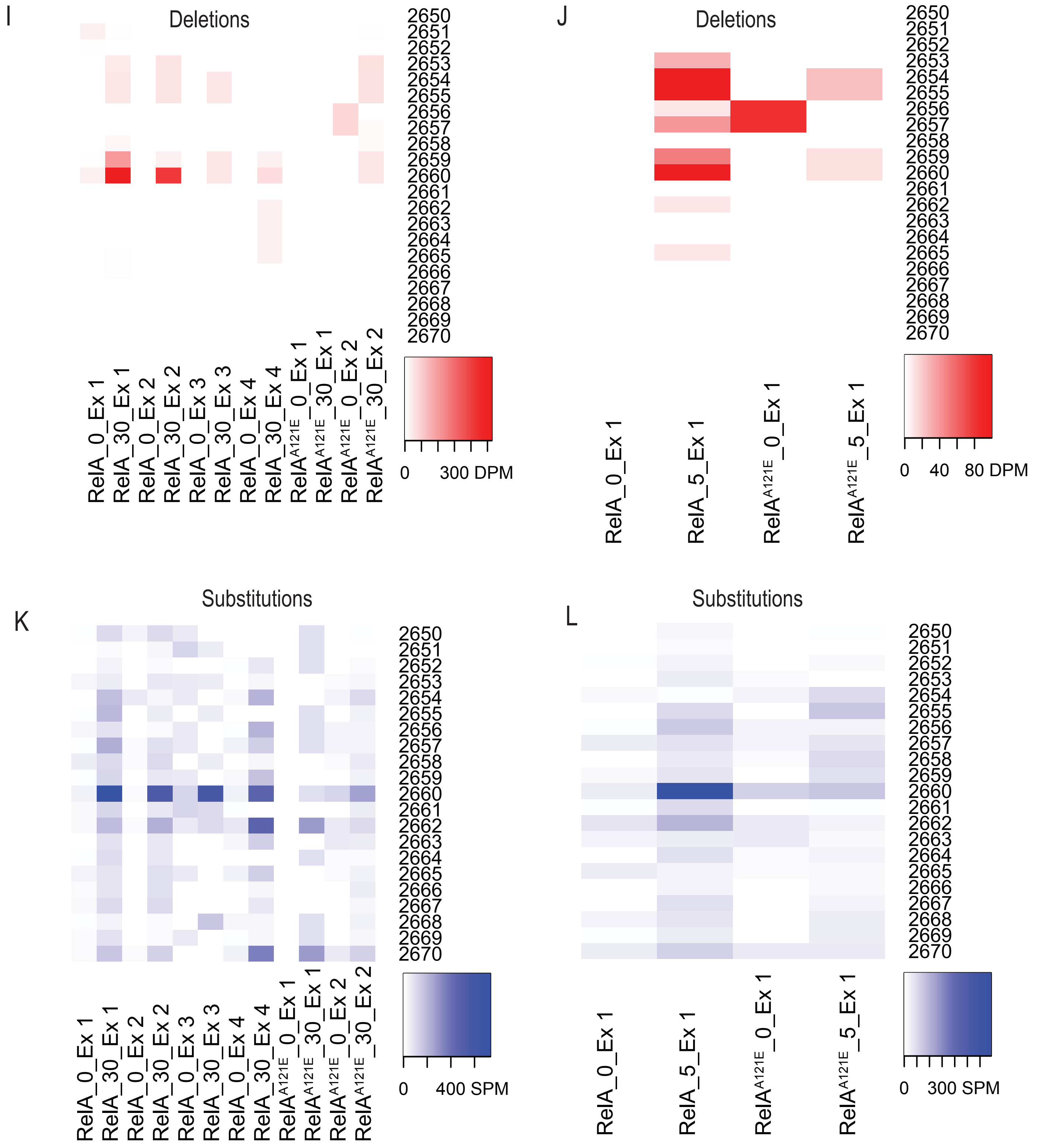
I
Deletions
RelA_0_Ex 1
RelA_30_Ex 1
RelA_0_Ex 2
RelA_30_Ex 2
RelA_0_Ex 3
RelA_30_Ex 3
RelA_0_Ex 4
RelA_30_Ex 4
RelA^A121E_0_Ex 1
RelA^A121E_30_Ex 1
RelA^A121E_0_Ex 2
RelA^A121E_30_Ex 2
0
300 DPM
J
Deletions
RelA_0_Ex 1
RelA_5_Ex 1
RelA^A121E_0_Ex 1
RelA^A121E_5_Ex 1
0
40
80 DPM
K
Substitutions
0
400 SPM
L
Substitutions
0
300 SPM

Fig. S4 - Page 3

M

Deletions

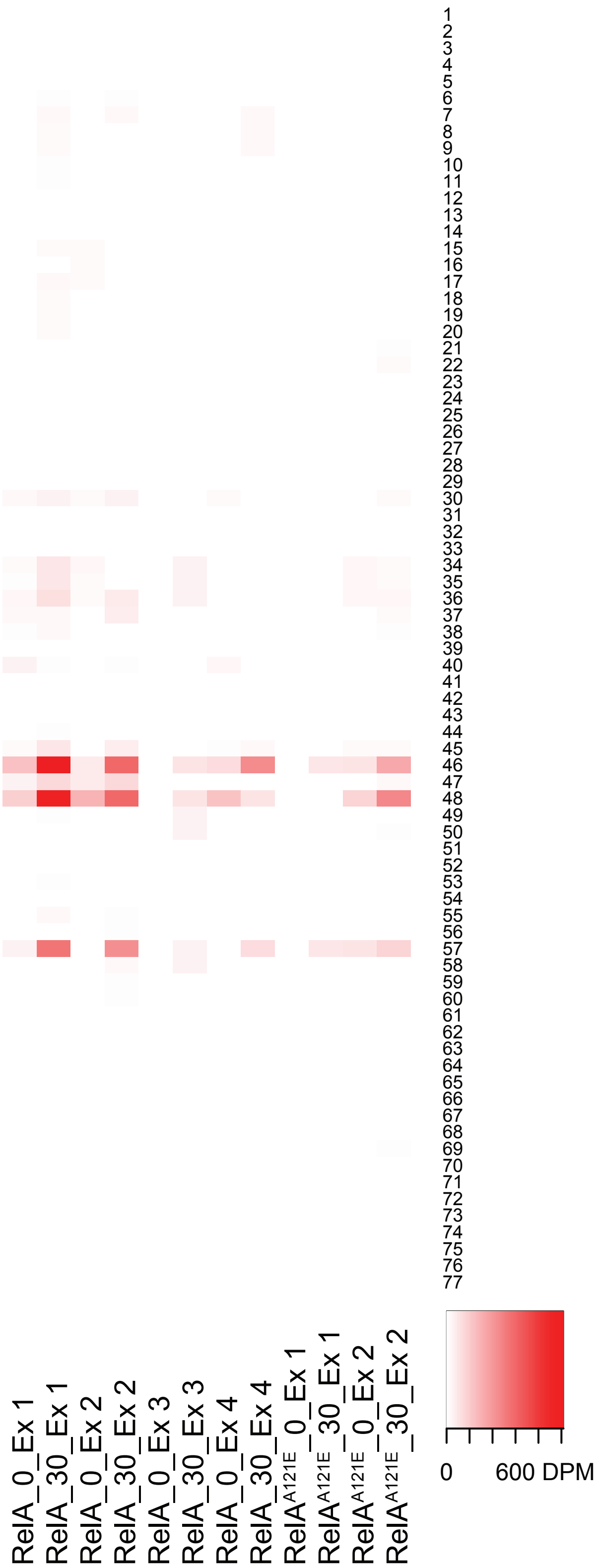

N

Substitutions

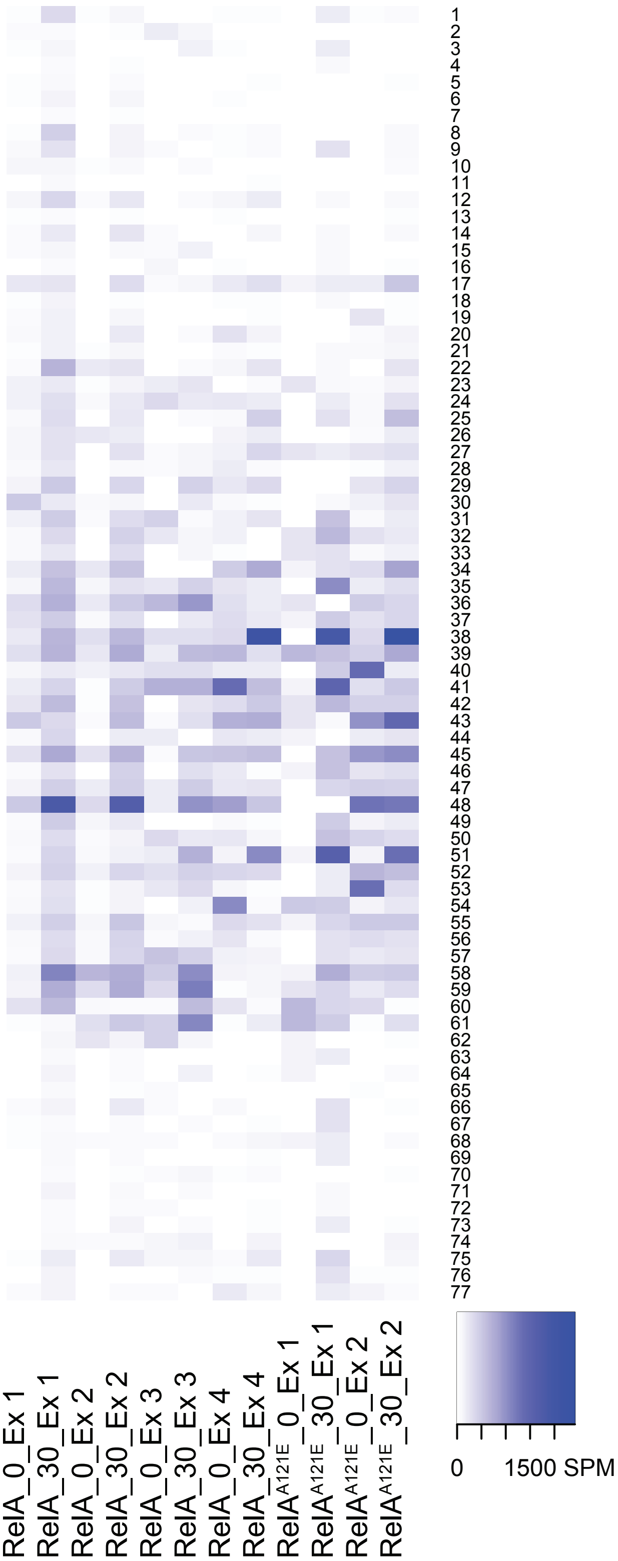

**Supplementary Fig. S4** Interaction-site mapping by cDNA mutational analysis. RT-mutations in cDNA read sequences obtained with CRAC analysis of MG1655 encoding *relA::HTF* or *relA*$^{A121E}$*::HTF* before (0) and after isoleucine starvation (5 or 30 minutes). **A)** and **B)** shows Deletions Per Million (DPM in red) cDNA reads mapped in the A-Site Finger (ASF) of 23S rRNA (nt 880-900) after 30 minutes or 5 minutes of starvation. **C)** and **D)** shows Substitutions Per Million (SPM in blue) mapped to the ASF. **E)** and **F)** shows DPM in cDNA reads mapped to Helix 15 of 16S rRNA (nt 355-375) after 30 minutes or 5 minutes of starvation. **G)** and **H)** shows SPM mapped to H15. **I)** and **J)** shows DPM in cDNA reads mapped to Sarcin-Ricin Loop of 23S rRNA (nt 2650-2670) after 30 minutes or 5 minutes of starvation. **K)** and **L)** shows SPM mapped to SRL. **N)** and **M)** shows deletions or substitutions per million in reads aligning to isoleucine tRNA (*ileT*) after 30 minutes of starvation.
